# Automatic decomposition of electrophysiological data into distinct non-sinusoidal oscillatory modes

**DOI:** 10.1101/2021.07.06.451245

**Authors:** Marco S. Fabus, Andrew J. Quinn, Catherine E. Warnaby, Mark W. Woolrich

## Abstract

Neurophysiological signals are often noisy, non-sinusoidal, and consist of transient bursts. Extraction and analysis of oscillatory features (such as waveform shape and cross-frequency coupling) in such datasets remains difficult. This limits our understanding of brain dynamics and its functional importance. Here, we develop Iterated Masking Empirical Mode Decomposition (itEMD), a method designed to decompose noisy and transient single channel data into relevant oscillatory modes in a flexible, fully data-driven way without the need for manual tuning. Based on Empirical Mode Decomposition (EMD), this technique can extract single-cycle waveform dynamics through phase-aligned instantaneous frequency. We test our method by extensive simulations across different noise, sparsity, and non-sinusoidality conditions. We find itEMD significantly improves the separation of data into distinct non-sinusoidal oscillatory components and robustly reproduces waveform shape across a wide range of relevant parameters. We further validate the technique on multi-modal, multi-species electrophysiological data. Our itEMD extracts known rat hippocampal theta waveform asymmetry and identifies subject-specific human occipital alpha without any prior assumptions about the frequencies contained in the signal. Notably, it does so with significantly less mode mixing compared to existing EMD-based methods. By reducing mode mixing and simplifying interpretation of EMD results, itEMD will enable new analyses into functional roles of neural signals in behaviour and disease.

**New & Noteworthy:** We introduce a novel, data-driven method to identify oscillations in neural recordings. This approach is based on Empirical Mode Decomposition and reduces mixing of components, one of its main problems. The technique is validated and compared with existing methods using simulations and real data. We show our method better extracts oscillations and their properties in highly noisy and non-sinusoidal datasets.

**Running Head:** Decomposition of data into non-sinusoidal oscillatory modes

## INTRODUCTION

The synchronised activity of neuronal populations can be observed in dynamic oscillations recorded in electrophysiology (1, 2). These oscillations are often visible in raw data traces but are challenging to isolate in an objective, data-driven manner. Methods for signal isolation must contend with signals being obscured by noise or by simultaneous oscillations at different frequencies. Neuronal oscillations are often non-sinusoidal and change over time, which leads to ambiguities in standard analyses based on the Fourier transform (3, 4). These dynamic and non-sinusoidal features are of growing importance in electrophysiological research but remain difficult to analyse using existing methods (1, 5–9). As such, there is a pressing need for data-driven methods that can isolate oscillations from noisy time-series whilst preserving their non-sinusoidal features.

Empirical Mode Decomposition (EMD) (10) is able to provide a different perspective on analysing transient oscillations. It offers a radically different approach to signal separation based on a flexible, local, and data-driven decomposition with weaker assumptions about stationarity and linearity of the signal. Single channel data is decomposed by a sifting process into Intrinsic Mode Functions (IMFs) based on finding successively slower extrema. Unlike Fourier or Wavelet methods, EMD does not a-priori assume the shape of its functions. It is therefore believed IMFs can capture non-sinusoidal oscillations and may better reflect the underlying processes in physical and physiological signals (3, 10, 11). This can especially aid analyses sensitive to waveform shape, such as calculations of phase and cross-frequency coupling (4, 12).

The original EMD algorithm can in theory produce arbitrarily shaped IMFs, but in noisy neural signals it struggles with signal intermittency and high non-sinusoidality. In the presence of transient oscillatory bursts, the sifting process may detect extrema on different time scales at different times. This is referred to as *mode mixing*. It presents a major challenge in analysis and interpretation of IMFs (13, 14). This is especially the case in analysis of brain signals, where transient states are common and have functional significance (5, 15–17). Furthermore, in the presence of pure Gaussian fractional noise, EMD has been shown to act as a dyadic filter bank (18, 19). This means that for highly noisy signals, EMD tends to produce IMFs with fixed bandwidths rather than adapting to capture signals present in the data, further complicating analysis.

Various improvements to the sifting process have been proposed to make EMD more applicable to real-world data (20–27). A unifying characteristic of the existing approaches is to inject a secondary signal into the data to alter the extrema distribution and overcome mode mixing. *Noise-assisted methods*, as exemplified by Ensemble EMD (EEMD) (20, 22), use white noise as the injected signal. This reduces mode mixing due to signal intermittency. However, the use of noise can limit IMF bandwidth, possibly making mode mixing worse. *Masking methods* inject sinusoids into the data before sifting (21, 24). With a suitable mask, this technique can recover non-sinusoidal waveforms and/or intermittent bursts in presence of noise. However, the frequency of masking signals that should be used is often not known a-priori. Mask optimisation can become an arduous manual process, prohibiting generalisability and introducing uncertainty on analysis outcomes. This is exacerbated by the presence of high noise and non-sinusoidal signals near dyadic boundaries, where a small change in the masking signal frequency may dramatically alter the quality of resulting IMFs. Mask frequency selection can be done semi-automatically by choosing an initial frequency based on the number of zero-crossings in the first IMF and dividing this successively by two for later IMFs (21). If the approximate frequency content of the signal is known, then mask frequencies may be directly selected to isolate the specific components of interest (11). Though effective, the semi-automatic method is relatively inflexible, and the direct specification method can be manually intensive to validate.

In this paper, we introduce iterated masking EMD (itEMD), a novel sifting technique that builds on the masking method. This method retains all the advantages of using a masking signal whilst being more generalisable and automated. We compare itEMD with existing methods using simulations and multi-species, multi-modal experimental data, and discuss its range of applicability and limitations.

In simulations, we have focused on the three areas important to the analysis of neural signals, as mentioned above: noise, sparsity, and waveform shape distortion. All three of these cause major issues for EMD and limit its applicability to neurophysiology. Here we show that itEMD performs significantly better than existing methods in cases with highly noisy and strongly non-sinusoidal signals, where our technique significantly reduces mode mixing and accurately extracts oscillations and their shape. We further validate the technique by analysing oscillations in rat local field potential (LFP) data and human magnetoencephalography (MEG) recordings. We show that itEMD reproduces the well-known hippocampal theta waveform shape better than existing techniques and successfully reconstructs occipital alpha peak frequency with no a-priori information about mask frequencies. Furthermore, itEMD significantly reduced mode mixing in both datasets studied.

## MATERIALS AND METHODS

### Data and Code Availability Statement

All figures and analysis in this paper can be freely replicated with Python code available at https://gitlab.com/marcoFabus/fabus2021_itemd. Hippocampal LFP data is available from the CRCNS platform (https://crcns.org/data-sets/hc/hc-3) and human MEG data is available from the Cam-CAN archive (https://camcan-archive.mrc-cbu.cam.ac.uk/dataaccess/) (28, 29). Analyses were carried out in Python 3.9.4, building on the open-source EMD package (v0.4.0), available with tutorials at https://emd.readthedocs.io (30). Underlying dependency packages were numpy (31), scipy (32), and statsmodels (33) for computation and matplotlib for visualisation.

### EMD Algorithms

Empirical Mode Decomposition decomposes a signal x(t) into a finite number of Intrinsic Mode Functions (IMFs) c_i_ with a *sifting* algorithm (10). The IMFs are constructed to have locally symmetric upper and lower envelopes with no peaks below zero or troughs above zero. A smooth signal with these features is well-behaved during instantaneous frequency analysis, allowing for a full description of non-sinusoidal waveform shape (11).

Ensemble EMD (20) is typical of a class of noise-assisted sifting methods. An ensemble of N sift processes is created, each with different white noise injected. The final IMFs are computed as the average across this ensemble. The goal is to exhaust all possible sifting solutions, leaving only persistent real signals. However, due to a finite size of the ensemble, IMFs may contain unwanted residual noise unless further improvements are introduced (26, 27). Furthermore, due to the stochastic nature of white noise, signals of interest might shift between modes across the ensemble, leading to some mode mixing in the final result. Finally, the use of noise reinforces the dyadic filtering behaviour of EMD. This means any signal near dyadic boundaries is likely to be split between modes. This effect is especially pronounced for non-sinusoidal signals which change in instantaneous frequency, making waveform shape analysis difficult as they become smeared over multiple IMFs.

Masked EMD (21) works by injecting a masking signal s_i_(t) into signal x(t) before sifting. This reduces mode mixing by making the sift ignore signal content slower than the frequency of the masking signal. The masking signal is introduced uniformly across n_p_ phases at each step to further minimise mode mixing (22). The IMFs c_i_ are thus calculated with the following algorithm:

1. Construct a masking signal s_i_(t).
2. Perform EMD on x_k_ = x(t) + s_i,k_(t + φ_k_), where φ_k_ = 2π(k-1) / n_p_, obtaining IMFs c_i,k_(t).
3. Compute the final IMF as c_i_(t) = 1/n_p_ ∑c_i,k_.
4. Compute the residue r_i_(t) = x(t) – c_i_(t).
5. Set x(t) = r_i_(t) and repeat 1-4 with the next masking signal to extract the next IMF.

This technique permits analysis with intermittent bursts and non-sinusoidal oscillations (Figure 1). EMD is locally adaptive, and as such fast bursts get mixed with slower activity when bursts are not present. With a mask, any signal content with frequencies much lower than the masking frequency will be ignored by the sift in that iteration and is replaced by the mask. The mask is finally removed allowing us to recover intermittent activity correctly. In presence of noise, EMD also acts as a dyadic filter (18, 19). This means non-sinusoidal oscillations are often split across multiple IMFs. With a suitable mask, the bandwidth of modes can be adapted and more of the waveform shape recovered.

**Figure 1:**
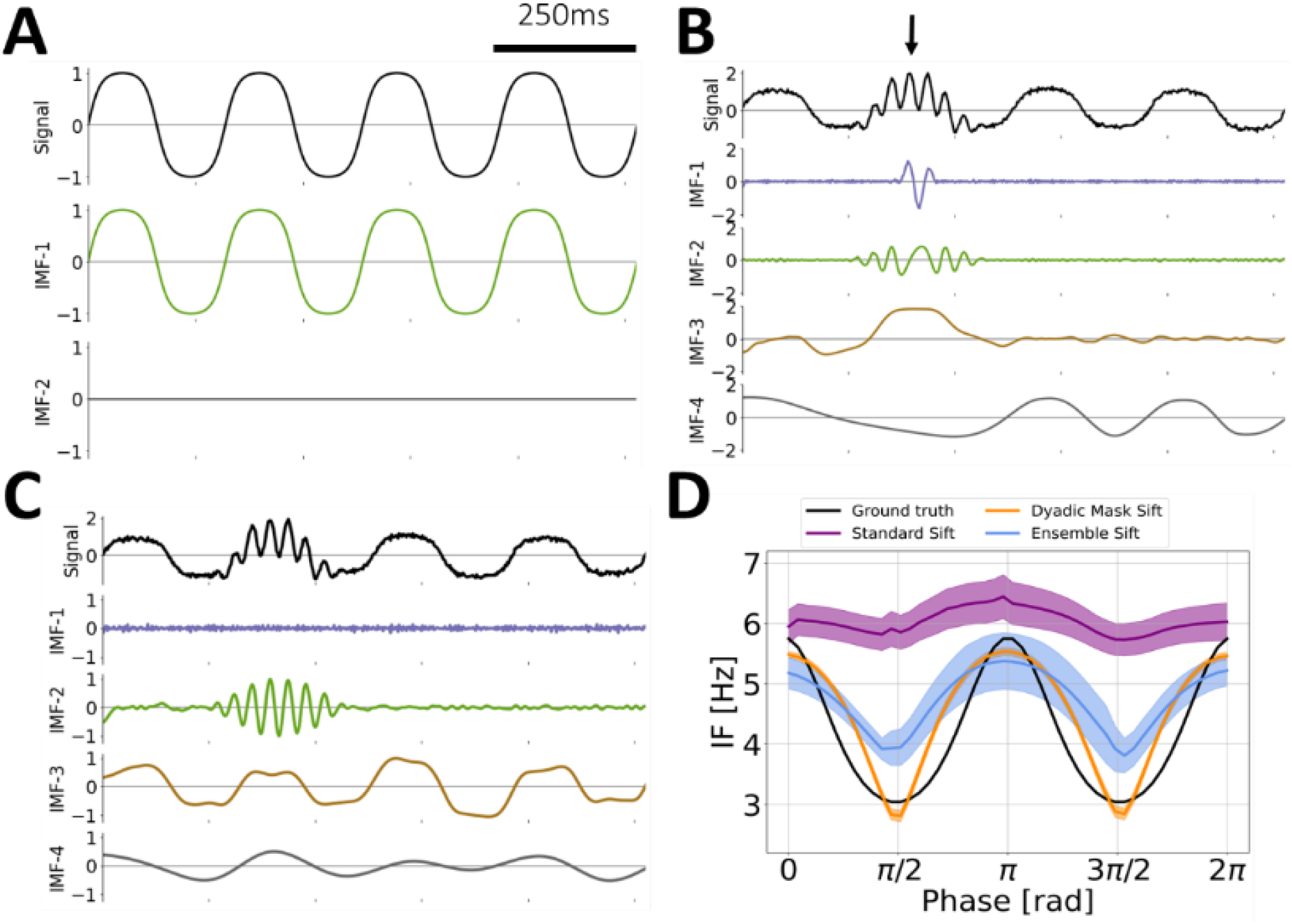
Limitations of EMD. (A) Standard EMD sifting applied on a pure 4Hz iterated sine function. With no noise, EMD can accurately identify an Intrinsic Mode Function (IMF) that represents the non-sinusoidal signal. (B) In presence of white noise and a 30Hz burst (arrow), standard EMD shows heavy mode mixing. (C) EMD with an appropriate dyadic mask sift will recover most of the iterated sine (IMF-3) and the intermittent burst (IMF-2) signals. (D) Masked EMD and Ensemble EMD can better reconstruct non-sinusoidal wave shape in signal with low noise unlike standard EMD. The figure shows the phase-aligned instantaneous frequency calculated from 100 runs of (B) and (C). Mean ± standard error (shaded) shown.

The choice of masking signals remains an area of active research. The original paper by Deering and Kaiser suggested the first mask frequency to be the energy-weighted mean of instantaneous frequency obtained from the first IMF found by ordinary EMD, with subsequent mask frequencies chosen as f_0_ divided by powers of 2 to account for the dyadic nature of EMD (21). Other approaches have included computing the mask from zero crossings of the first IMF of a standard sift and purely dyadic masks (24). However, the choice of optimal masks remains a manual process in many cases. This requires experience and may introduce subjective bias (11, 21, 25, 35).

### Iterated Masking EMD (itEMD)

As seen above, noise-assisted and masking approaches to EMD sifting improve mode mixing in some cases, but mode mixing may still be present to complicate further analysis. Mask choice in noisy datasets is complicated, especially with signal frequencies near dyadic boundaries. Iterated masking solves this problem by finding and using an adaptive, data-driven mask.

In science, it is common to rely on intuition to guide study of complex dynamical systems (36). Consider then a simple example where there is a signal burst x(t) with some base frequency f_sig_ and possible deviation around it due to non-sinusoidality. Take as a start the masked EMD process with a single mask of frequency f_mask^(0)^_. A good choice of frequency would be near f_mask_ = f_sig_, as this would extract most of x(t) into one IMF, resulting in noise reduction and allowing for a simple IMF interpretation. This is because adding a mask at f_mask_ = f_sig_ forces the IMF to ignore any spectral content below ∼0.67* f_mask_ (25).

In real data however, f_sig_ is often unknown. Assume then f_mask^(0)^_ is chosen with little to no knowledge of the system frequency f_sig_. After applying masked EMD, the resulting IMF will contain a part of the burst with some noise or other signal mixed in. However, its instantaneous frequency will be f_sig_ for sections of the IMF attributable to the signal. Assuming signal amplitude is distinguishable from noise in this IMF, the amplitude-weighted instantaneous frequency mean (AW-IFM) will be closer to the desired f_sig_ than f_mask^(0)^_. Thus, if we use this AW-IFM as the masking frequency for the next iteration f_mask_(^1^), the resulting mask sift IMF will be even closer to the optimal IMF. This is the case both if f_mask^(0)^_ is greater and smaller than f_sig_, as both lead to mode mixing. Following this reasoning, the natural equilibrium of this iteration process is when f_mask_ = f_sig_, and we can apply this approach to a signal consisting of multiple signal frequencies and noise. This leads us to the following algorithm:

1. Choose an initial set of mask frequencies m = {f_0_}.
2. Perform masked EMD to obtain IMFs.
3. Find the instantaneous frequency (IF) for each IMF using the Hilbert transform.
4. Compute the amplitude-weighted average of each IMF’s IF and set m_i_ = AW-IFM.
5. Repeat 2-4 until a stopping criterion ∑ is reached.

Here the stopping criterion was chosen such that the relative difference between current and previous mask frequencies is small, i.e. (m_i_ –m_i-1_) / – m_i-1_ < ∑. Instantaneous frequency averaging was weighted by the square of instantaneous amplitude for a given IMF, i.e. by instantaneous power. Mask frequencies were initialised by the dyadic masking technique, though it was found that itEMD is not sensitive to mask changes and can rapidly identify correct IMFs even with a random initial mask (Figure 2). Due to rapid convergence (<10 iterations in most cases), itEMD is computationally comparable to existing techniques including ensemble EMD and uniform phase EMD, each of which requires repeated sifting (22). More formally, the computational complexity of itEMD is T = 41n_iter_ * n_s_ * n_p_ * n log_2_(n) for n_iter_ iterations, n_s_ sifting steps, n_p_ mask phases, and data length n.

**Figure 2:**
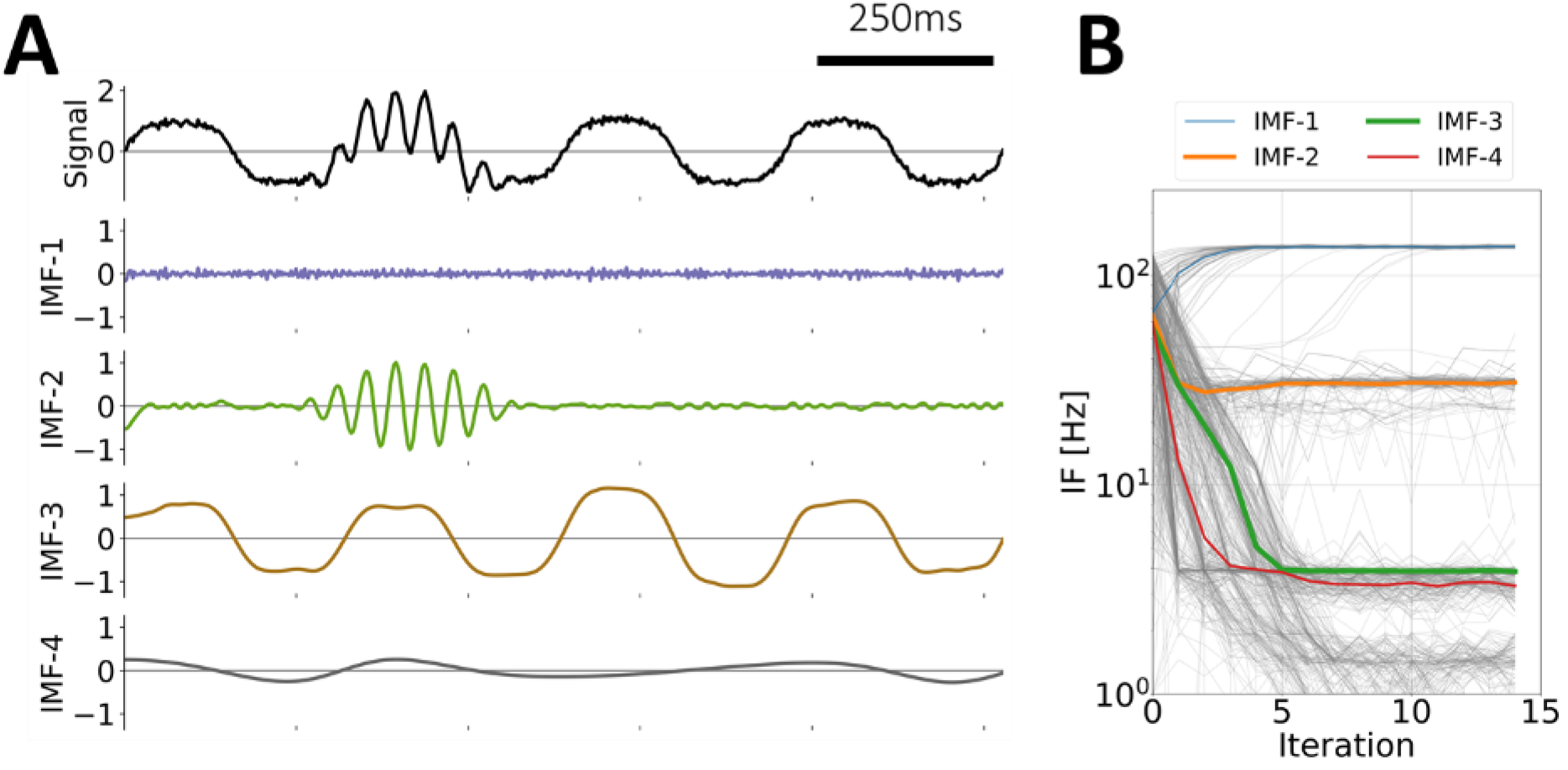
Iterated Masking EMD (itEMD) on simulated data. (A) Example endpoint of itEMD, i.e. IMFs from last of 15 iterations on the same signal as in Figure 1B. (B) itEMD convergence to equilibrium starting with a mask drawn randomly from a uniform distribution of 1-128Hz. Thin gray lines are all 100 individual runs, coloured lines are median trajectories with line thickness scaled to maximum value of that IMF. itEMD converges rapidly, adapts mask frequencies to the signal content, and correctly finds both the non-sinusoidal 4Hz oscillation and intermittent 30Hz burst with no prior knowledge about the frequencies contained in the signal.

### Simulations

We ran simulations to compare the performance of itEMD to existing sifting methods. Simulations were performed along three dimensions that are important to analysis of neural signals: noise, sparsity, and waveform shape distortion (non-sinusoidality). These were chosen as they are all common features of neurophysiological data which cause issues for extracting neural oscillations. In standard EMD, they result in mode mixing and prohibit accurate representation of waveform shape and robust interpretation of identified modes.

All noise and frequency distortion simulations were 10s long and sampled at 512Hz with signal amplitude normalised to 1. In each simulation, IMFs were computed using three different methods that are used to address mode mixing: Dyadic Mask Sift, Ensemble Sift, and our novel iterated masking EMD (itEMD). Dyadic Mask Sift utilised a single set of masking frequencies. The first was computed from zero-crossings of first IMF obtained by a standard sift and subsequent masking frequencies were divided by powers of 2. Masks were applied with 4 phases uniformly spread across 0 to 2π following Wang et al. (2016) (22). Ensemble sift was run with 4 noise realisations and ensemble noise standard deviation of 0.2. The novel itEMD was run on top of the masked EMD implementation with a stopping criterion ∑ = 0.1 and maximum number of iterations N_max_ = 15. In all simulations, number of IMFs was limited to 6 and the sifting threshold was 1e-08. After finding IMFs, individual cycles were found from jumps in the instantaneous phase found by the amplitude-normalised Hilbert transform. Each set of simulations (noise, distortion, sparsity) was repeated N=100 times with the mean +/- standard error results presented.

Waveform shape was quantified by computing the average phase-aligned instantaneous frequency (IF) across cycles (11). IF measures how an oscillation speeds up or slows down within a cycle. It is computed as the time derivative of the instantaneous phase. IF was phase-aligned to correct for differences in timing and duration between cycles and allow for comparisons at each phase point. It can intuitively be understood as fitting a sinusoid with frequency that of the instantaneous frequency at each time point, capturing shape deviations away from a sinusoid with a constant frequency (37, 38). Within-cycle IF variability is thus a measure of how non-sinusoidal each cycle is.

Performance of each method was assessed by two methods. The first was finding Pearson correlation between reconstructed phase-aligned instantaneous frequency (proxy for waveform shape) and its ground truth. The second was computing the Pseudo-Mode Splitting Index (PMSI) introduced by Wang et al. (2018) (22). PMSI estimates the degree of mode mixing between two adjacent IMFs by computing the normalised dot product between them:

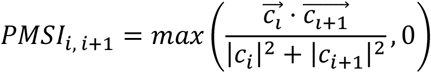

Orthogonal, well-separated modes with no mode mixing thus have PMSI=0. Fully split modes have PMSI=0.5. This index was chosen as it can be applied to both simulated and real data and is easy to interpret. For simulations with a known ground truth, we took the IMF of interest as the one with mean instantaneous frequency closest to that of the ground truth and calculated PMSI as the sum of PMSIs with the above and below IMF.

### Noise Simulations

For analysing noise-dependent properties, white noise was created using the numpy.random.normal Python module with zero mean and standard deviation σ (also equal to its root-mean-square, RMS). White noise was chosen as method performance on it is independent of signal frequency. This is because white noise has equal power throughout the frequency spectrum. For simulations of neurophysiological data, signals with brown noise were also considered (see Supplementary Information). For this set of simulations, white noise RMS σ was varied between σ=0.05 and σ=3 in 100 uniformly spaced steps. Waveform shape distortion was held constant at FD=68% (see below).

### Waveform Shape Distortion Simulations

For analysing waveform shape, signal was simulated as an iterated sine function, i.e. sin(sin(…sin(2*pi*f_0_*x))) iterated N_sin_ times with f_0_=4Hz. This function was chosen because i) it is easy to manipulate its non-sinusoidal distortion by increasing N_sin_, ii) it is well-understood analytically (39), iii) it has been used before in context of EEG time-frequency analysis (40), and iv) it has a well-behaved instantaneous frequency by satisfying conditions outlined in Huang et.al. (1998) (10). It also qualitatively captures parts of waveform shape of the sensorimotor mu oscillation and slow oscillations in depth EEG recordings by its ‘flat top’ structure (7, 41). The base frequency of 4Hz was chosen as it is physiologically plausible in the delta range and was near a Nyquist boundary, where current EMD sifting methods may have issues. Its non-sinusoidality was captured by a frequency distortion metric FD defined by

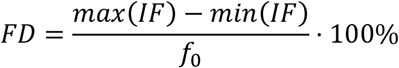

A signal with FD=0% is a pure sinusoid and FD=100% indicates a waveform with IF range equal to that of the original frequency, i.e. 4±2Hz. An example waveform can be seen in Figure 1 (FD=68%). In this set of simulations, frequency distortion was varied between FD=18% and 101% by repeating simulations with iterated sine order varying from N_sin_=1 to N_sin_=18. White noise RMS was held constant at σ = 1.

### Signal Intermittency Simulations

For analysing effects of signal intermittency on itEMD performance, we simulated bursts of different length in a 25s segment of data. Sparsity was measured as the number of individual oscillations in the burst. The number of cycles in the burst was varied from 5 to 95 in 100 steps. Noise RMS was kept constant at σ_noise_ = 1 and distortion at FD=68% (8^th^ order iterated sine).

Statistical testing was done using one-sided Welch’s t-test corrected for multiple comparisons using Bonferroni’s method unless otherwise specified (42).

## Experimental Data

### Rat Local Field Potential (LFP) Data

To validate our method with well-described hippocampal theta oscillations, we used a publicly available data set of Long-Evans rats (43, 44). The full 1000s local field potentials (LFP) recording from rat EC-013 sampled at 1250Hz was used for analysis. The electrode analysed was implanted in the hippocampal region CA1. EMD cycle analysis was the same as during simulations. In short, three types of sifting methods were compared: dyadic masking sift with zero-crossing initialisation, ensemble sift, and the novel itEMD. The recording was split into 20 segments of 50s duration before sifting. For itEMD (as in simulations), the stopping criterion was set at ∑=0.1, the maximum number of iterations was N_max_=15, the mask was weighted by squared instantaneous amplitude, and the iteration process was initialised by the zero-crossing dyadic mask result. Instantaneous phase, frequency, and amplitude were computed from the IMFs using the amplitude-normalised Hilbert transform with an instantaneous phase smoothing window of N=5 timepoints. The theta IMF was chosen as that whose average instantaneous frequency was closest to the Fourier spectral theta peak estimated using Welch’s method (peak in 4-8Hz, function scipy.signal.welch, 8s segment length / 0.125Hz resolution). Cycles were computed from jumps in the wrapped instantaneous phase. To discard noisy cycles, only cycles with monotonic instantaneous phase, instantaneous amplitude above the 50^th^ percentile, and instantaneous frequency below 16Hz were used for further analysis. Cycles were phase-aligned with N=48 phase points and the shape was represented by the mean of the phase-aligned instantaneous frequency. To compare mode mixing, the PMSI (see above) was also computed as the sum of PMSIs of the theta IMF with the IMF above and below it in frequency.

### Human Magnetoencephalography (MEG) Data

Ten resting state MEG recordings were randomly chosen from the CamCAN project (https://www.cam-can.org/) (28, 29). The participants were randomly chosen from the project (mean age 43.5 years, range 18-79, 6 female). The maxfilter processed data were downloaded from the server and converted into SPM12 format for further analysis using the OHBA Software Library (OSL; https://ohba-analysis.github.io/osl-docs/). Each dataset was down-sampled to 400Hz and bandpass filtered between 0.1 and 125Hz. Two notch filters were applied at 48-52Hz and 98-102Hz to attenuate line noise. Physiological artefacts were removed from the data using Independent Components Analysis. 62 components were computed from the sensor space data and artefactual components identified by correlation with EOG and ECG recordings. Any independent component with a correlation greater 0.35 with either the EOG or ECG was considered artefactual and removed from the analysis. This resulted in two to four components removed from each dataset. EMD analyses proceeded with the cleaned MEG data from a single gradiometer MEG2112 over midline occipital cortex. Each recording was about 10 minutes (median 565s, range 562s-656s). The power spectrum of the whole recording was estimated using Welch’s method (function scipy.signal.welch, 8s segment length / 0.125Hz resolution). The frequency of the spectral alpha peak was then extracted in the 8-12Hz range as a local maximum (function scipy.signal.find_peaks). For itEMD analysis, each recording was segmented into 10 parts of the same length (median segment length 56.2s). EMD was performed with the mask sift, ensemble sift, and itEMD. Sift parameters were identical to those used for the rat LFP analysis above. The IMF representing alpha oscillations was chosen as the one whose mean instantaneous frequency was closest to the alpha peak frequency. Subjects were excluded if no alpha peak was present (one subject). After extraction of cycles from the Hilbert-computed instantaneous phase jumps, only those with instantaneous frequency between 7Hz and 14Hz and instantaneous amplitude above the 50^th^ percentile were kept. For further analysis, cycles were phase-aligned to N=48 uniformly spaced phase points between 0 and 2π and the mean across cycles was computed for each subject. To evaluate mode mixing, the PMSI for each sifting method was also calculated.

## RESULTS

### Simulations

Iterated masking sift (itEMD) rapidly converged on signal in presence of noise and intermittency. An initial ten second data segment with a 30Hz transient burst, a 4Hz non-sinusoidal oscillation, and low white noise was first simulated (Figure 2). The iteration process was started with a set of six random masks drawn uniformly from 1-128Hz. Despite this initial complete lack of knowledge about the signal, itEMD correctly recovered the non-sinusoidal waveform and the beta-frequency burst. The iteration process converged with noise in IMF-1, the 30Hz beta burst in IMF-2, and non-sinusoidal 4Hz signal in IMF-3. Subsequently, convergence was determined automatically. The convergence criterion was set to the mask stabilising within 10% between iterations with a maximum number of iterations of 15 (see Methods). Further simulations were initialised with the zero-crossing masked sift results for faster convergence. All simulations were repeated with N=100 different noise realisations.

### Influence of noise

We wanted to establish how different sifting methods reconstruct waveform shape in presence of noise (Figure 3). Ten seconds of a 4Hz non-sinusoidal iterated sine signal with white noise was simulated. The standard deviation of zero-mean white noise (root-mean-square of noise, RMS noise) was varied as the shape of the wave remained constant with 8^th^ order iterated sine (frequency distortion FD=68%, see Methods). Iterated masking was compared to existing dyadic masking and ensemble sifting techniques.

**Figure 3:**
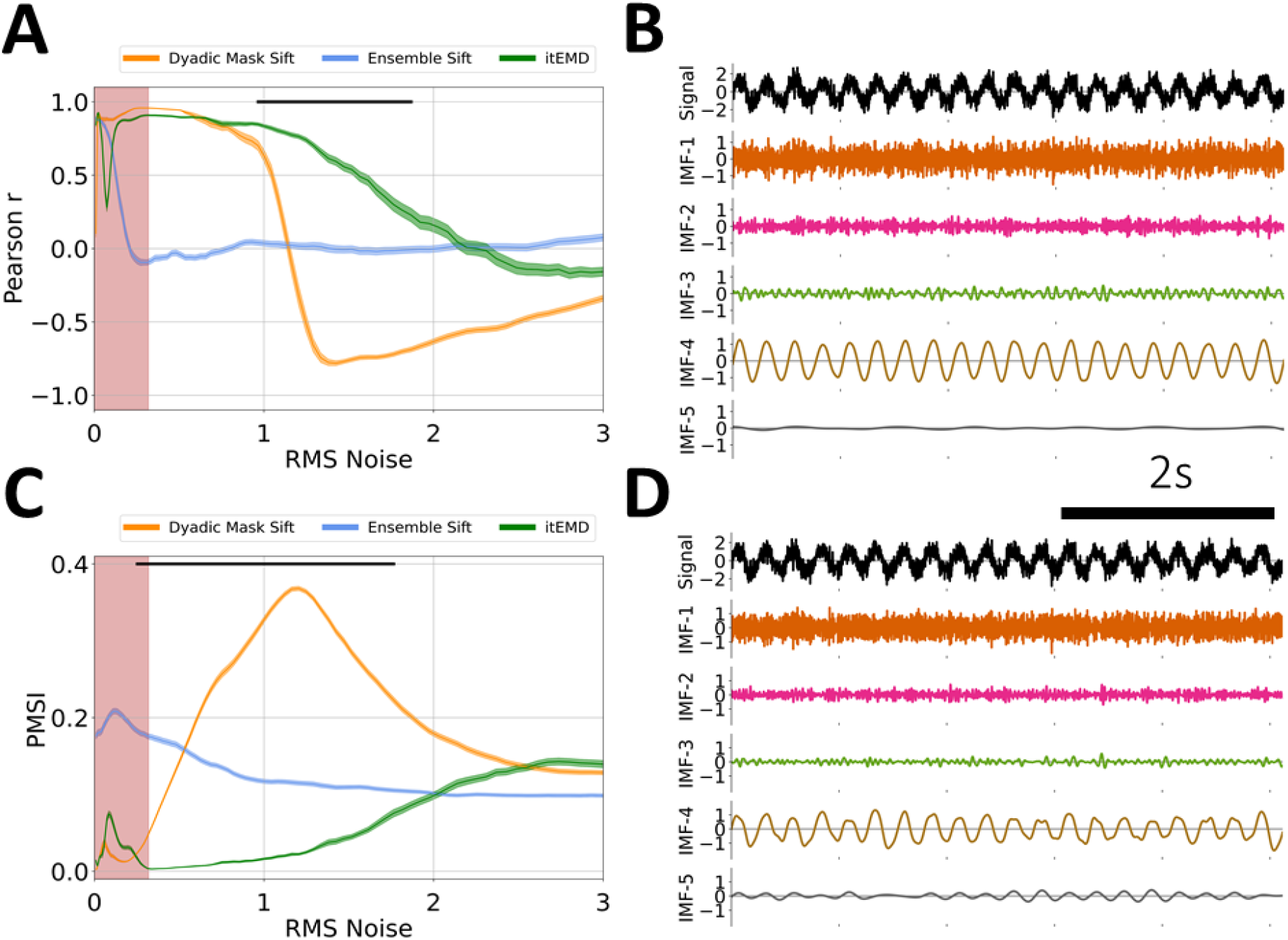
Influence of noise on EMD performance on simulated data. (A) Pearson correlation coefficient between reconstructed and ground truth instantaneous frequency against increasing white noise amplitude. (B) Example 5s of itEMD sift results for σ_noise_ = 0.5, FD=68%. Iterated sine function is captured by IMF-4. (C) Pseudo-Mode Splitting Index (PMSI) against white noise amplitude, with higher PMSI values indicating higher mode mixing. Mean ± standard error across N=100 noise realisations shown. Black line indicates regions where itEMD performs significantly better than the best of the other techniques with P<0.01 (multiple comparisons Bonferroni corrected). Red shaded region shows noise levels where itEMD reached maximum number of iterations in >20% of noise realisations. The novel itEMD performs significantly better for highly noisy data in the region between σ_noise_ ≈ 1 and σ_noise_ ≈ 2 with reduced mode mixing and accurate waveshape reconstruction. (D) Example dyadic mask sift results for σ_noise_ = 0.5, FD=68%. IMF-4 shows significantly more mode splitting than itEMD results.

First, we measured performance by computing Pearson correlation of reconstructed and ground truth waveform shape measured by the instantaneous frequency (Figure 3A).

For low to medium noise amplitudes (σ_noise_ ⪅ 1 with normalised signal amplitude of one), existing techniques were sufficient to represent the waveform shape well. Ensemble-sift reconstructed waveform shape had a high correlation with the ground truth shape with r > 0.75, but its performance quickly deteriorated past σ_noise_ = 0.1. Dyadic mask sift had poor shape reconstruction for no noise but performed well from σ_noise_ = 0.1 onward. The novel iterated masking performed well except for a dip in performance around ultra-low noise below σ_noise_ = 0.1. This was found to be the level where noise RMS is equal to the amplitude of one of the higher signal harmonics. As such, this harmonic was sometimes included in the IMF of interest and sometimes not, depending on exact noise details. This introduced stochastic mode mixing (see also Supplementary Information and Figure S1). Noise levels in neurophysiological data are seldom this low. However, we were able to automatically identify these pathological cases because the mask did not reach a stable equilibrium and the maximum number of iterations was reached (red shading).

For high noise amplitudes (1 ⪅ σ_noise_ ⪅ 2), the new itEMD significantly outperformed existing techniques. Ensemble sift produced sine waves and failed to capture non-sinusoidal waveform behaviour (correlation near zero). Dyadic mask sift suffered from mode mixing, with the waveform split across IMF-4 and IMF-5 (Figure 1D). Because of this, dyadic masking failed to accurately reconstruct waveform shape, especially above σ_noise_ = 1. In contrast, itEMD accurately isolated the signal even at high noise levels (Figure 3B). Its reconstructed shape correlated with the ground truth significantly better than the existing techniques with P < 0.01 (Bonferroni corrected across 100 noise levels) across the high noise range 1 ⪅ σ_noise_ ⪅ 2.

In the region of very high noise with σ_noise_ > 2, all methods behaved as dyadic filters and failed to capture waveform shape. This is because higher Fourier harmonics encoding shape details became submerged in noise, making it impossible to recover the non-sinusoidal shape.

We also evaluated mode mixing performance by computing the PMSI (Pseudo – Mode Splitting Index, Figure 3C), a mode mixing metric previously used in the literature (22). A high PMSI value indicates severe mode mixing and poor sift.

For low noise amplitudes (σ_noise_ ⪅ 0.3), the dyadic mask sift produced the least amount of mode mixing (lowest PMSI). In this region, itEMD was again susceptible to stochastic mode mixing due to noise levels matching higher harmonics, increasing the PMSI. Ensemble sift had the most mode mixing in this region.

For medium to high noise (0.3 ⪅ σ_noise_ ⪅ 2), itEMD had significantly less mode mixing than existing techniques (lowest PMSI P < 0.01, Bonferroni-corrected). Ensemble sift had a largely unchanging amount of mode mixing, suggesting it was driven by the added noise. Dyadic masking had the most mode mixing in this region.

All three methods had similar PMSI in the very high noise region with σ_noise_ > 2 due to inherent dyadic filtering behaviour of EMD.

Neurophysiological signals typically show auto-correlated 1/f noise (also termed aperiodic activity or fractal noise) (45). To verify our technique works with 1/f noise simulations, we re-ran all the main analyses with brown noise (Supplementary Information S3). As with white noise, itEMD outperformed existing techniques over a wide range of parameters.

Finally, we compared mask frequency stability across itEMD iterations for a mode known to have signal (IMF-4) and a pure noise mode (IMF-5). The mask frequency was significantly less variable when signal was present (Supplementary Information S5, Figure S5).

### Influence of frequency distortion (non-sinusoidality)

Highly non-sinusoidal waveforms have been observed across a variety of neural data (see Introduction). As such, we compared existing techniques and itEMD performance in data with progressively more waveform distortion. Ten seconds of a 4Hz non-sinusoidal iterated sine signal with white noise of standard deviation σ_noise_ = 1 was simulated. Frequency distortion was varied by iterating the sine function between 1 and 18 times. Each frequency distortion level was simulated with N=100 different noise realisations. Performance was again compared using Pearson correlation to ground truth shape and the PMSI.

Iterated masking performed significantly better than existing methods for highly non-sinusoidal signals (Figure 4, Bonferroni-corrected P<0.01 for lowest PMSI and higher Pearson r). Dyadic mask shape correlation with ground truth was not significantly different from itEMD for FD < 50%, but severe mode mixing was present. This meant the average frequency and amplitude were poorly reconstructed. At this noise level, ensemble sift behaved as a dyadic filter and completely failed to capture waveform shape. It produced a sinusoid at the dyadic boundary of f=4Hz with no non-sinusoidality. itEMD performance also improved with increasing frequency distortion. This is due to higher frequency harmonics increasing in magnitude with more shape distortion, allowing better convergence of itEMD.

**Figure 4:**
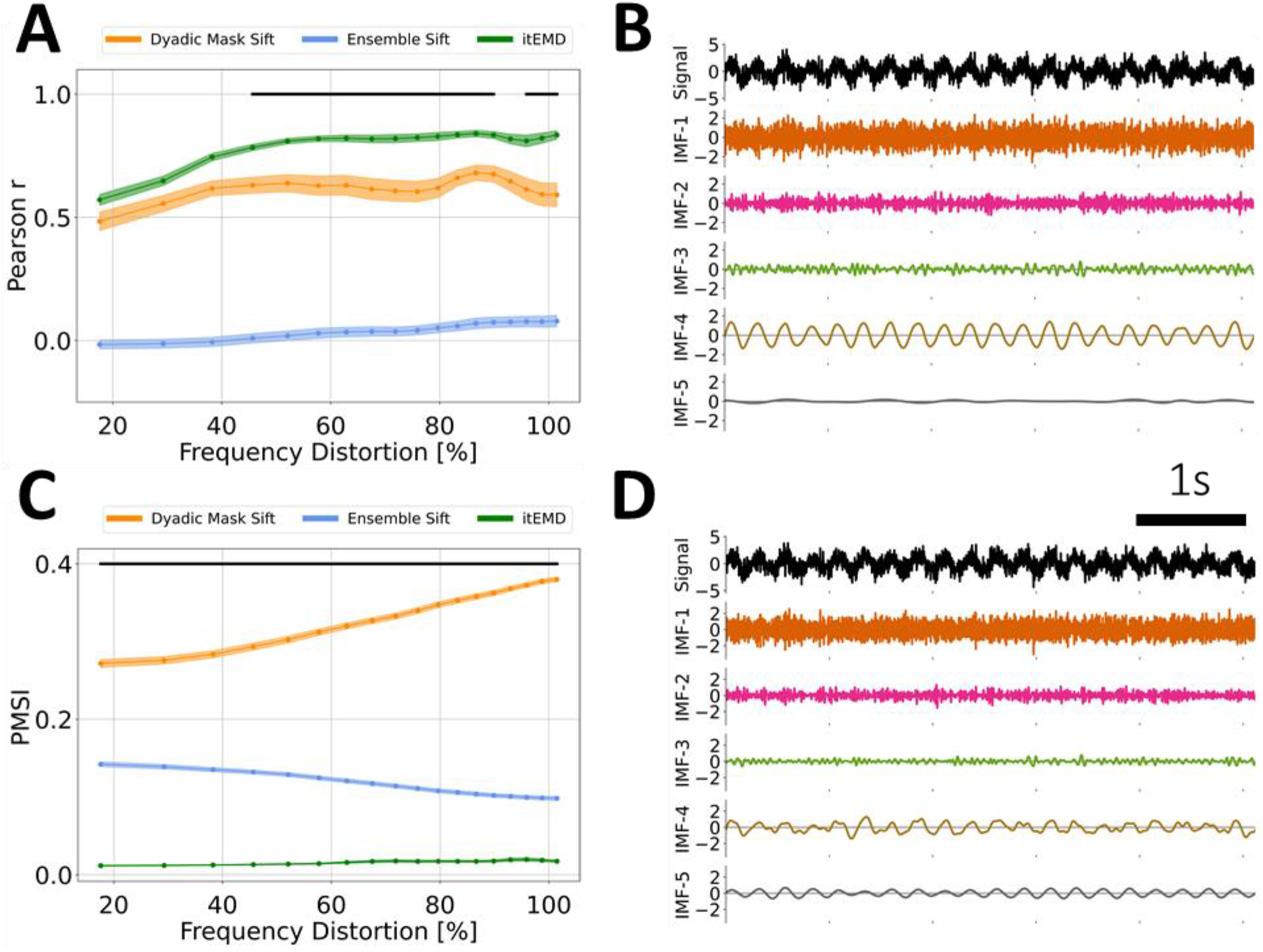
Influence of frequency distortion on EMD performance in simulated data. (A) Pearson correlation coefficient between reconstructed and ground truth instantaneous frequency against increasing frequency distortion. (B) Example 5s of itEMD sift results for σ_noise_ = 1, FD=80%. (C) Pseudo-Mode Splitting Index (PMSI) against frequency distortion, with higher PMSI values indicating higher mode mixing. Mean ± standard error across N=100 noise realisations (shaded) shown. Black line indicates regions where itEMD performs significantly better than the best of the other techniques with P<0.01 (multiple comparisons Bonferroni corrected). The novel itEMD performs significantly better for highly non-sinusoidal data in the region FD > 50% with reduced mode mixing and accurate waveshape reconstruction. (D) Example 5s of dyadic mask sift results for σ_noise_ = 1, FD=80%. IMF-4 shows significantly more mode mixing with existing methods than with our

### Reconstructed Waveform

Next, we looked at individual IMFs and reconstructed waveforms and their instantaneous frequency (Figure 5). As expected from the Pearson r and PMSI results in figure 4, itEMD best reconstructed a highly non-sinusoidal waveform in the presence of noise. A noise level of σ_noise_ = 0.1 and 4^th^ order iterated sine were chosen as they are qualitatively similar to experimental LFP and MEG recordings analysed (cross-reference Figure 7). The ensemble sift was able to capture most of the non-sinusoidality but suffered from heavy mode mixing (PMSI = 0.0943). The dyadic mask sift had slightly less mode mixing (PMSI = 0.0923) but failed to capture any non-sinusoidal waveform shape details. The novel iterated masking captured the waveform shape best with the least mode mixing (PMSI = 0.0003). However, the waveform was still not perfectly reconstructed. This was due to i) some of the harmonics encoding the finer details being lower in spectral density than the noise and ii) due to intrinsic finite bandwidth of EMD modes (see Supplementary Information and Figure S2). However, itEMD performed significantly better than the other techniques with the root-mean-square error to the ground truth instantaneous frequency being significantly lower (P=1.75e-9 vs dyadic mask, P=0.045 vs ensemble sift, Bonferroni-corrected).

**Figure 5:**
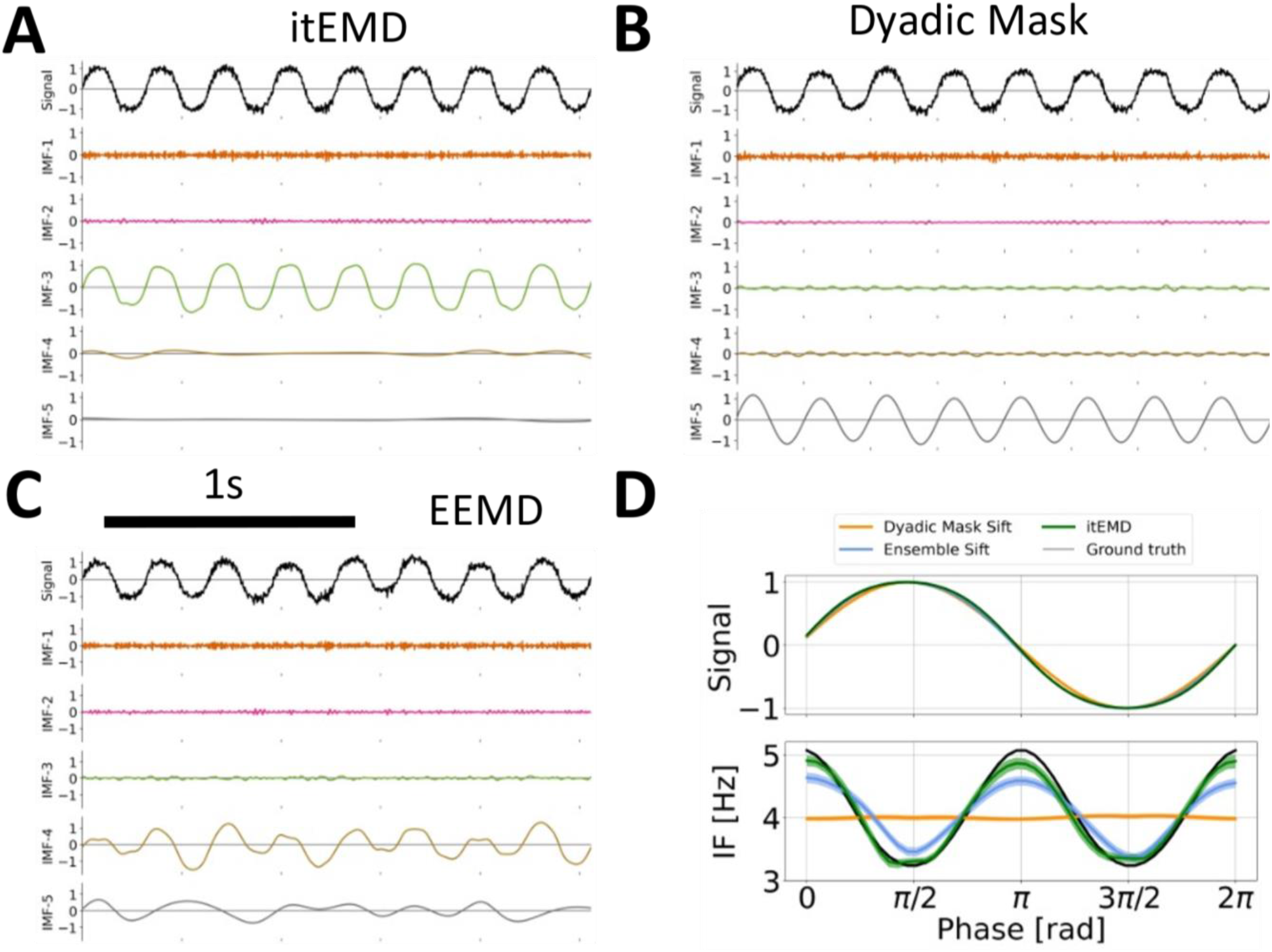
Example 2s of sifting results for 4^th^ order iterated sine with white noise σ_noise_ = 0.1 in simulated data. (A) IMFs for the novel itEMD. Iterated sine is in IMF-3 with very little mode mixing. (B) IMFs for dyadic mask sifting. Signal is split between IMF-3 and IMF-5. (C) IMFs for ensemble sifting. Iterated sine is mostly in IMF-4, but mode mixing is present. (D) Top: average reconstructed waveform, bottom: reconstructed phase-aligned instantaneous frequency (IF); mean (line) ± standard error across cycles (shaded) shown. Dyadic mask sift waveform (orange) fails to reconstruct non-sinusoidality. Ensemble sift recovers more shape detail but suffers from high mode mixing. itEMD is able to reconstruct more of the waveform shape than either existing method whilst lowering mode mixing.

### Influence of signal sparsity

Neural activity often consists of intermittent bursts (17). To test itEMD performance when signal is sparse, we simulated 25 seconds of zero-mean white noise with σ_noise_ = 1, to which we added a 4Hz non-sinusoidal 8^th^ order iterated sine signal with frequency distortion FD = 68% and variable length of 5-100 cycles. When reconstructing waveform shape of this burst, itEMD performed significantly better than either the dyadic mask sift, or the ensemble sift (Figure 6). Even in the presence of high noise and non-sinusoidality, itEMD was able to extract the burst and identify its waveform shape. Its correlation with ground truth waveform shape was significantly higher than the other methods for all burst lengths considered (Figure 6A, P < 0.01, Bonferroni corrected across number of cycles in the burst). Mode mixing measured by the PMSI was also significantly lower than with existing methods (Figure 6C, P < 0.01, Bonferroni corrected). Performance of itEMD improved as burst length increased. Overall, this demonstrates the potential benefits of itEMD when characterising transient bursts, which is increasingly used to describe oscillations in electrophysiological data (5).

**Figure 6:**
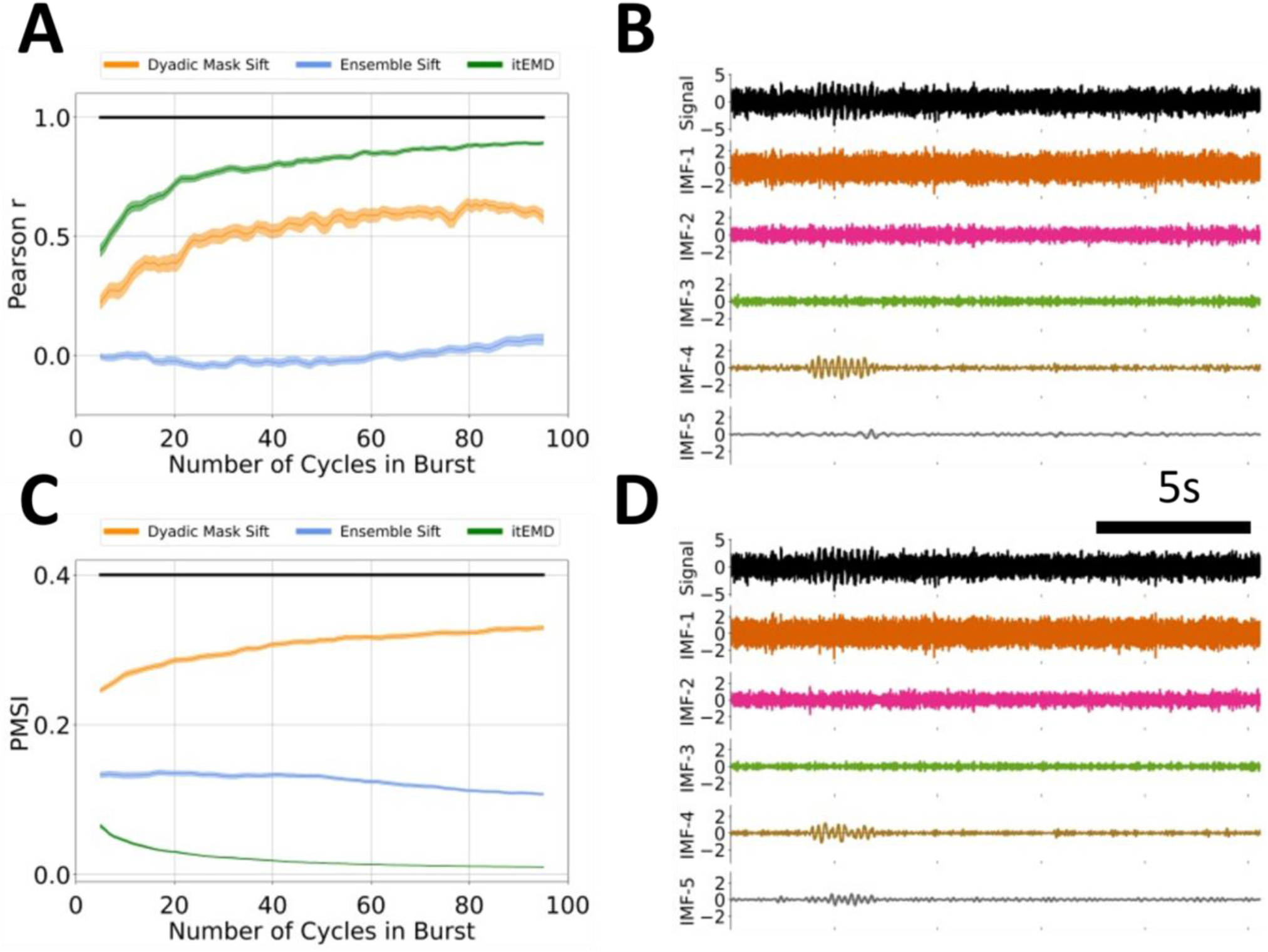
Influence of signal sparsity on EMD performance on simulated data. (A) Pearson correlation coefficient between reconstructed and ground truth instantaneous frequency against increasing burst length. (B) Example itEMD sift results for σ_noise_ = 1, FD=68%, 10 cycles. (C) Pseudo-Mode Splitting Index (PMSI) against burst length. Mean ± standard error across N=100 noise realisations (shaded) shown. Black line indicates regions where itEMD performs significantly better than the best existing technique with P<0.01 (multiple comparisons Bonferroni corrected). The novel itEMD performs significantly better for a wide range of burst durations. (D) Example dyadic mask sift results for σ_noise_ = 1, FD=60%, 10 cycles in burst. Iterated sine function is captured by IMF-4 and IMF-5 due to mode mixing.

## Application to experimental data

### Rat Local Field Potential (LFP)

We first validated our technique by applying it to the well-understood hippocampal theta signal in a 1000s recording of publicly available rat hippocampal LFP data. The recording was split into N=20 segments of 50s each. This theta oscillation has been previously observed to be non-sinusoidal with, on average, a faster ascending than descending edge (8, 46, 47). Our novel iterated masking EMD (itEMD) converged after N_iter_ = 6±1 iterations and extracted cycles with a wide instantaneous frequency sweep (Figure 7). It reproduced the known shape with a faster leading edge (leading edge frequency 7.87±0.02Hz, falling edge frequency 7.62±0.02Hz, mean ± SEM, P=5.9e-21 on a paired t-test across all cycles). In comparison to itEMD, existing ensemble and dyadic mask sifting failed to capture the high non-sinusoidality of this oscillation. Existing methods also suffered from higher mode mixing as measured by the PMSI (lowest PMSI for itEMD with Bonferroni-corrected P < 10^-6^). This was confirmed by visualising the Hilbert-Huang transforms, where theta IMF has the cleanest sweep for itEMD. This could allow for improved cross-frequency coupling analysis.

**Figure 7:**
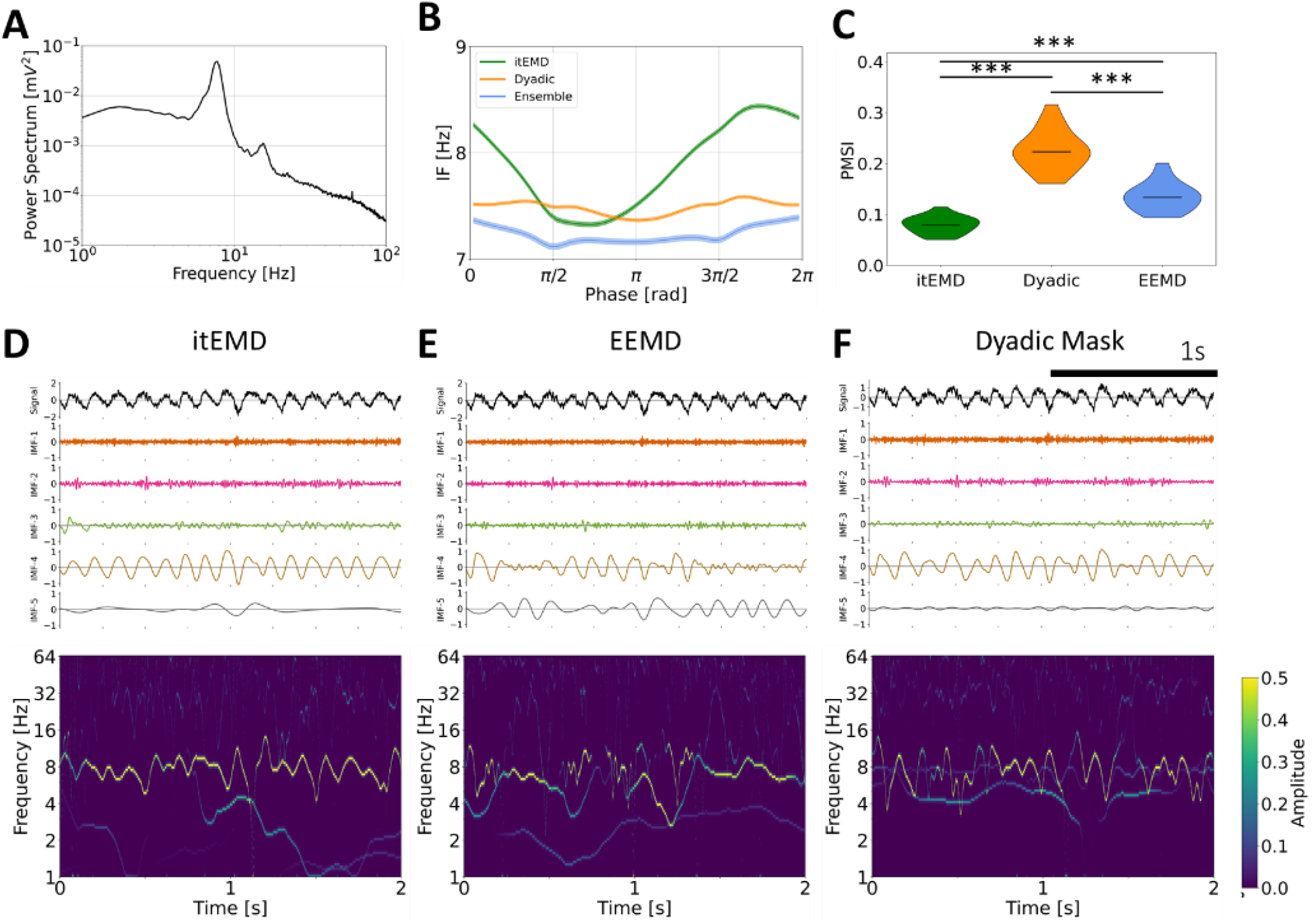
Rat hippocampal LFP results. (A) Power spectrum of the full recording showing a theta peak a harmonic. (B) Phase-aligned instantaneous frequency of theta cycles (mean ± standard error across all cycles shown). Existing methods including dyadic mask sift and ensemble sift fail to capture high non-sinusoidality of theta oscillations unlike itEMD. (C) Violin plots of the pseudo-mode splitting index (PMSI, a measure of mode mixing) across N=20 segments of the 1000s recording. Iterated masking had significantly lower PMSI than both dyadic mask sift and ensemble sift (P < 10^-6^, Bonferroni-corrected across methods). (D) Top - example itEMD sift results from two seconds of the LFP recording, bottom – Hilbert-Huang Transform (HHT) for the same region. Theta oscillations are well-captured by IMF-4 with minimal mode mixing. (E) Ensemble sift results from the same recording. Significant mode mixing is present complicating further analysis. (F) Dyadic mask sift results for the same recording. An artefactual low-frequency component is present.

### Human Magnetoencephalography (MEG)

For further validation, we analysed 10 minutes of occipital resting-state data from each of 10 subjects (Figure 8). One subject was excluded as their spectrum did not show an alpha peak. We found itEMD successfully and rapidly converged on the intermittent alpha oscillation around 10Hz (N_iter_ = 5±1 iterations across all subjects, mean ± standard deviation). Compared to Dyadic mask sift and ensemble sift, mode mixing measured by the PMSI was significantly lower (P = 6.0e-5 vs dyadic mask, P = 0.0033 vs ensemble sift, Bonferroni-corrected paired t-test across subjects).

**Figure 8:**
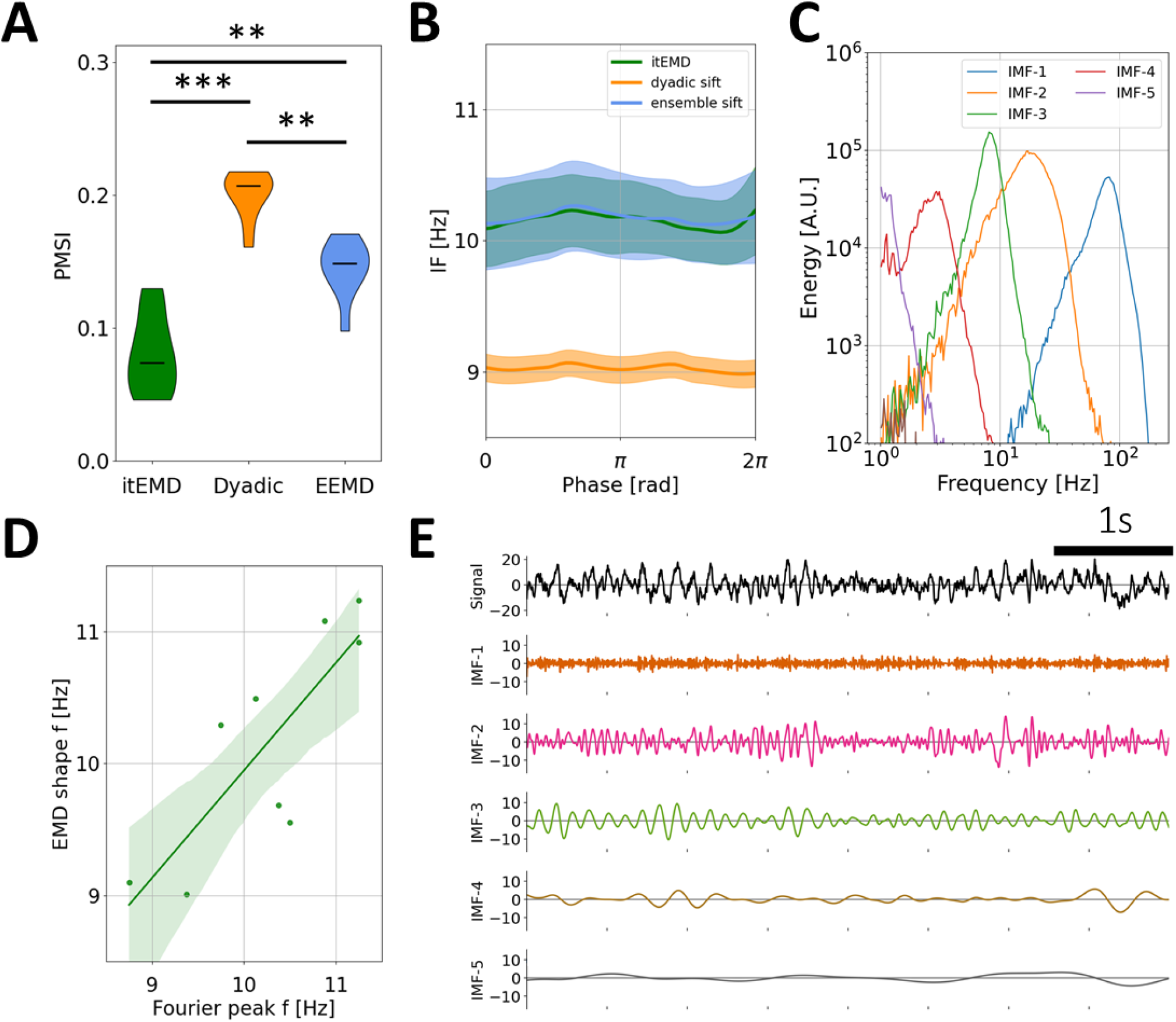
Human MEG occipital alpha results. (A) Group-level violin plots of the pseudo-mode splitting index (PMSI, a measure of mode mixing). Iterated masking had significantly lower PMSI than both dyadic mask sift (P = 6e-5), and ensemble sift (P = 0.0062, both Bonferroni-corrected). (B) Group-level phase-aligned instantaneous frequency (mean± standard error shaded). Both itEMD and EEMD detect the 10Hz occipital alpha oscillation with no significant non-sinusoidality. Masked sift fails to capture the oscillation well due to mode mixing. (C) 1D HHT for IMFs from an example subject. (D) Mean subject phase-aligned instantaneous frequency against peak alpha frequency from the Fourier power spectral density. Iterated masking linearly reproduces between-subject variability in alpha frequency (P=0.00739, F-test against constant null hypothesis). (E) Example 5s of raw itEMD sift results. Alpha oscillations are in IMF-3 with sharp features in IMF-2 and minimal mode mixing.

Alpha peak frequency is known to vary between people, conditions, and changes with age (48, 49). EMD has the advantage of representing spectral components with fewer assumptions about frequency bands, unlike traditional Fourier analyses focusing on the 8-12Hz power for example. As such, it may be well-suited to represent these inter- and intra-individual spectral differences. Variability in alpha peak frequency was also seen in data analysed here. Moreover, itEMD was able to replicate this result, with the mean subject phase-aligned instantaneous frequency found to be linearly related to spectral peak frequency (Figure 8D, F = 13.89, P = 0.00739). Recordings also showed the presence of beta-band elements around 20Hz in IMF-2, which were also present in the Fourier spectrograms (Figure 8E and Supplementary Figure S5-C).

## DISCUSSION

In this paper, we introduced a novel way of performing Empirical Mode Decomposition (EMD) called iterated masking EMD (itEMD). This technique is capable of robustly decomposing signals into spectral components in presence of noisy, sparse, and highly non-sinusoidal oscillations. In itEMD, masking signals are introduced at frequencies identified by an iterative, data-driven process in order to solve the mode mixing observed with other EMD methods. We demonstrated the utility of this sifting technique in comparison with existing solutions to the mode mixing problem, especially in highly noisy and non-sinusoidal signals.

We validated the method using rat LFP and human MEG recordings. We found rat hippocampal theta to be highly non-sinusoidal with a faster ascending edge as previously reported (8, 46, 47). Intermittent human occipital alpha was found to be nearly sinusoidal with a between-subject variable peak frequency around 10Hz. Mode mixing was found to be significantly lower when using itEMD compared to ensemble and dyadic mask sifting in these neurophysiological recordings. Iterated masking EMD has the potential to enable more wide-spread use of EMD in neurophysiology and shed light on single-cycle dynamics across a wide range of modalities and conditions. It automates the selection of mask frequencies and can thus enable a wide range of analyses about bursts of neural activity, genuine cross-frequency coupling, and analysis of neural phase.

We performed extensive simulations across a wide range of noise, sparsity, and non-sinusoidality (frequency distortion) parameters. In our validation of itEMD using simulated data, we used white noise. We focused on white noise because in this case results become independent of simulated signal frequency and hence ought to be applicable to any common noise structure. However, our technique is just as applicable to different noise structures (e.g. brown noise, Supplementary Information S3).

The stability of the mask equilibrium found by itEMD was found to be higher for modes containing genuine signal (Supplementary Figure S5). It was observed that the equilibrium was well-defined for IMFs displaying consistent oscillations with between-iteration mask frequency changes of less than 3% and rapid convergence (<10 iterations). This makes sense as itEMD found the equilibrium where IMF frequency matched mask frequency, as described in the Methods. Conversely, IMFs containing mostly noise had highly variable mask frequencies changing by more than 5% between iterations with no well-defined equilibrium. This could be used to assess whether an oscillation is present in any given segment of data in addition to existing ways of doing this such as amplitude and period consistency (35).

### Comparison with existing analysis methods

As itEMD is an EMD-based technique, this paper focused on direct comparisons with other EMD sifting techniques. As mentioned, a key reason why EMD has scarcely been applied to neurophysiological data is mode mixing (13, 14). When oscillations of interest are present across multiple IMFs, their interpretation and further analysis is made much more difficult. Iterated masking EMD significantly reduced mode mixing (measured by the PMSI as per previous literature before (22)) whilst still being able to reconstruct non-sinusoidal oscillations in the signal. Compared with the existing masked sift, itEMD has the advantage of being fully data-driven and avoiding manual mask optimisation.

Non-EMD based analysis methods may complement itEMD. Traditionally, analysis has been done by calculating the Hilbert transform on narrowband filtered data (8). This works well if frequencies of interest are defined a priori. However, it poses limitations on how non-sinusoidal oscillations can be and does not allow for large between-subject variabilities. Furthermore, the use of Fourier filters may introduce bias into the analysis (10, 50). More recently, methods based on detecting phase control points (peaks, troughs, etc) have been developed (35). These provide important summary statistics for cycles, such as peak-trough asymmetry and rise-decay asymmetry. EMD-based analysis describes the shape with phase-aligned instantaneous frequency without restricting analysis to certain phase points. The cycle-by-cycle approach could thus be cross-validated by itEMD detecting asymmetry around the phase points used for its statistics. Finally, additional novel algorithms for extracting summary waveforms for a whole recording have been developed (51, 52). Unlike itEMD and the techniques described above, these are however not sensitive to changes in waveform within a recording.

### Limitations when using itEMD

Although itEMD represents a significant step towards extracting non-sinusoidal neural oscillations in a data-driven way, there may be situations where other techniques are more appropriate. Here we draw attention to a few cases where this may be the case.

Iterated masking EMD works to capture more waveform shape details by adapting the bandwidth of an IMF to include more signal from higher frequency harmonics (Supplementary Information S2). However, together with capturing more shape details, this also increases the amount of noise in the IMF. If oscillations being studied are not expected to be non-sinusoidal, or such features are not of interest, other methods (such as a carefully designed manual mask for masked sift, or Fourier techniques) may be better at boosting signal-to-noise ratio. Indeed, it was found that at high noise levels, the SNR boost from itEMD is lower than other sifting methods (Supplementary Figure S2).

As with most EMD-based algorithms, itEMD also has a fundamental limitation in how large a single IMF bandwidth can be. It has been previously shown that the original EMD algorithm differentiates between a single amplitude-modulated tone and two separate tones based on their rates of extrema and amplitude ratios (53). This means that when shape-encoding harmonics are much higher in frequency than the base frequency (approximately *af^2^ > 1* for amplitude ratio *a* and frequency ratio *f* of base to harmonic as explained in (53)), itEMD will tend to treat these as two separate oscillations. If they need to be treated as a single oscillation, researchers should use masks specifically designed for overcoming this limit (25) or non-EMD based waveform analysis techniques (e.g. cycle-by-cycle analysis (35)). It is still a matter of debate as to when higher-frequency signals constitute harmonics as opposed to genuine separate oscillations (54, 55). This issue is important, as harmonics can cause spurious cross-frequency coupling if not accounted for properly, and ongoing research in our group is attempting to clarify this issue. Finally, in high levels of noise, high-frequency harmonics may be below the noise level (Supplementary Information S1). When this happens, itEMD will only partially reconstruct the waveform shape. This is because EMD works locally by finding extrema, and if they are dominated by noise on a given scale, EMD will not be able to identify signal. Our technique shares this limitation with all existing EMD-based analysis methods. However, even at high noise levels when this happens, our technique still had significantly lower mode mixing and preserved the correlation between ground truth and reconstructed instantaneous frequency.

itEMD was designed to handle sparse oscillations, but it may be necessary to adjust the amplitude weighting method if signals of interest are very sparse (<10% of a data segment being processed). This can be done by changing the weighting of instantaneous frequency when iterating. Here we used weighting by the square of instantaneous amplitude (IA^2^, instantaneous power) at each iteration. Higher powers of instantaneous amplitude may help if sparsity is preventing itEMD from converging on oscillations of interest. Conversely, weighting by lower powers may be appropriate if minute IA fluctuations are driving spurious results.

In this work, we defined iteration convergence when the relative mask change between iterations was under 10%. We also tried continuing for ten iterations after this point and averaging the resulting IMFs to verify the robustness of our threshold. It was qualitatively observed that only minimal changes occurred after the 10% convergence point. However, in datasets not studied here, it may be necessary to tune the convergence criterion or take the average of a few iterations after soft convergence around 10% depending on exact noise structure present.

When analysing our MEG data, we segmented the recordings into 10 parts of about a minute each. itEMD implicitly assumes that the mean frequency of oscillations of interest is not changing greatly and an equilibrium can be found. Hence, to allow for drifting in the spectral peak frequency segmented data was used. However, it was found alpha frequency did not vary significantly during this resting-state experiment and applying itEMD on full recordings yielded very similar results (see Supplementary Information S5). In general, we recommend segmenting the data if peak frequencies may shift over time (e.g. in drug induction or task data).

In summary, we have introduced a novel way to robustly extract oscillatory modes from neural recordings using iterated masking EMD. Our method has all the advantages of using EMD whilst resolving limitations of existing sifting techniques by significantly reducing the mode mixing and robustly capturing oscillations even in presence of noise, sparsity, and high non-sinusoidality. By validating it on extensive simulations and real multi-modal, multi-species data, we have demonstrated its potential to bring the full power of EMD into neurophysiology and help elucidate the role of dynamic neural oscillations in behaviour and disease.

## AUTHOR CONTRIBUTIONS

Conceptualisation: MSF.

Methodology: MSF, AJQ, KW, MWW.

Software: MSF, AJQ.

Formal analysis: MSF.

Writing Original Draft: MSF.

Writing Review & Edit: MSF, AJQ, KW, MWW.

Supervision: KW and MWW.

Funding Acquisition: KW and MWW.

## ACKNOWLEDGMENTS & FUNDING

This research was funded in part by the Wellcome Trust [Grant number 203139/Z/16/Z]. For the purpose of open access, the authors have applied a CC-BY public copyright licence to any Author Accepted Manuscript version arising from this submission.

Research was also supported by the NIHR Oxford Health Biomedical Research Centre, the Wellcome Trust (106183/Z/14/Z, 215573/Z/19/Z), the New Therapeutics in Alzheimer’s Diseases (NTAD) study supported by UK MRC and the Dementia Platform UK (RG94383/RG89702), an EU European Training Network grant (euSSN; 860563), and the MRC Development Pathway Funding Scheme (award reference MR/R006423/1).

Data collection and sharing for this project was provided by the Cambridge Centre for Ageing and Neuroscience (CamCAN). CamCAN funding was provided by the UK Biotechnology and Biological Sciences Research Council (grant number BB/H008217/1), together with support from the UK Medical Research Council and University of Cambridge, UK.

## SUPPLEMENTARY INFORMATION

### S1. Performance at very low noise levels

Iterated masking sift experienced a dip in performance for very low noise levels around σ=0.05. Mode mixing measured by the PMSI increased significantly and waveform shape reconstruction decreased in accuracy. This can be understood by examining the power spectrum around this point (Figure S1). For ultra-low noise levels, first four harmonics of the iterated sine function are well resolved, and most of the iterated sine function is in IMF-3. For higher noise at σ=0.1 at this level of frequency distortion, the fourth harmonic is barely detectable due to increased noise levels. Iterated sine is sifted mainly into IMF-4. In-between these noise levels (around σ=0.05), mode mixing occurs because the fourth harmonic is above noise for some cycles and below it for others. From an equivalent time-domain perspective, this process can be seen as whether the extrema on a given level are determined mainly by noise or by higher harmonics of the signal. Iterated masking assumes waveform shape details (i.e., higher Fourier harmonics) are well-resolved, and may experience worse performance when this is not the case. However, as mentioned in the Main Text, such cases were easy to automatically identify. The mask failed to converge at the 10% level and maximum number of iterations was reached.

**Figure S1:**
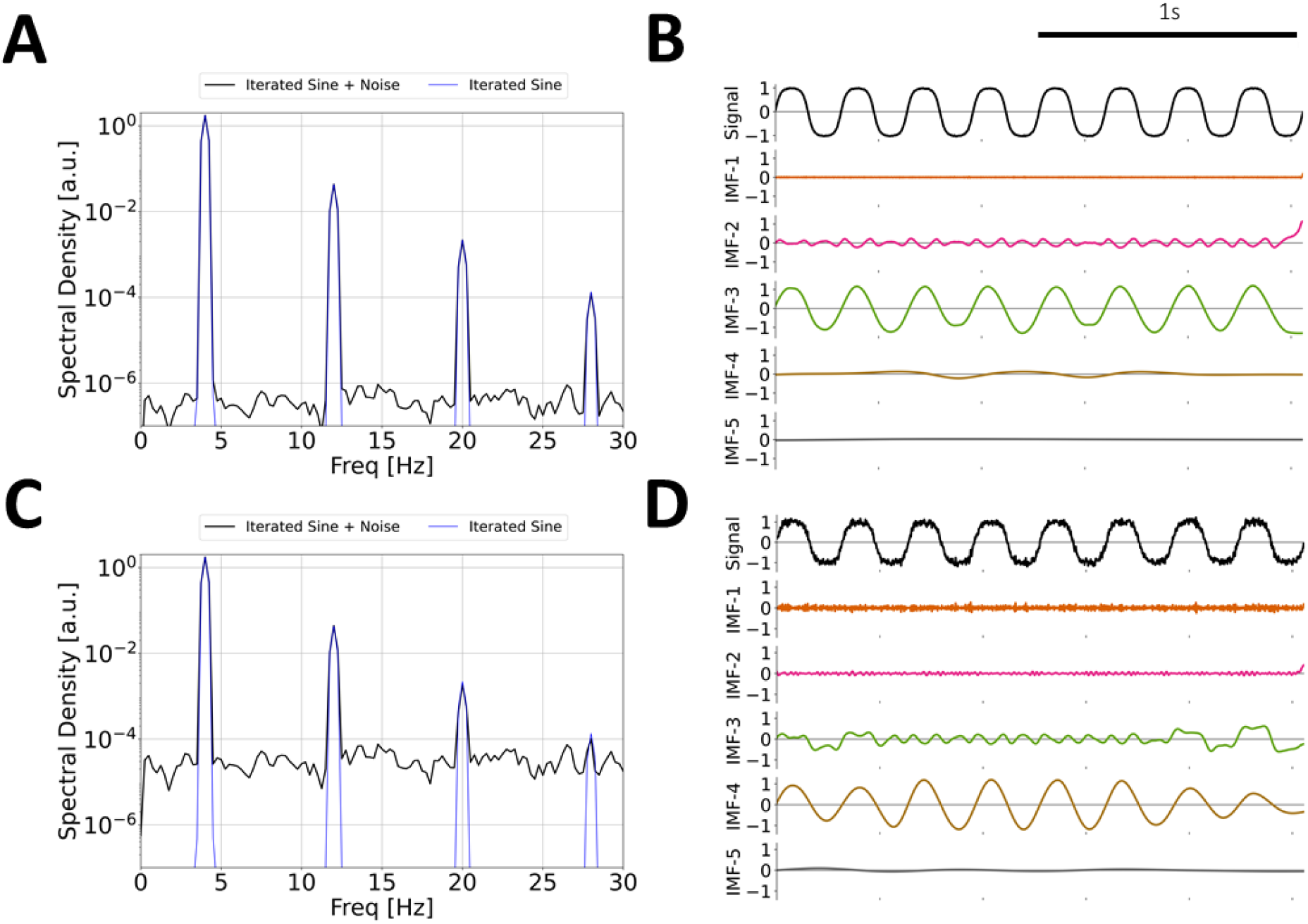
Performance of itEMD dips for very low noise levels around σ=0.05. (A) Power spectral density of data for ultra-low noise with σ=0.01 (black) and pure 8^th^ order iterated sine (blue). (B) Sift results for σ=0.01, with 8^th^ order iterated sine in IMF-3. (C) Power spectral density of data for noise with σ=0.1. (D) Sift results for σ=0.1, with 8^th^ order iterated sine in IMF-4 and some mode mixing.

### S2. Bandwidth limitations of EMD

As discussed in the Introduction, EMD acts as a dyadic filter bank for high noise datasets (18, 19). More precisely, this means sifting splits signal sampled at frequency *F_s_* into IMFs with bandwidth [*F*_*S*_/2^*N*^, *F*_*S*_/2^*N*+1^] for *N>0*. This is detrimental to analysis of neural signals because non-sinusoidal oscillations near dyadic boundaries are split between modes; EMD is similar to simply applying a series of Fourier filters. As such, IMFs with adaptive bandwidths covering signal harmonics / shape details are more desirable. To study itEMD bandwidth, ten seconds of 4Hz 8^th^ order iterated sine signal with varying zero-mean white noise were simulated. The spectral bandwidth was calculated as the frequency range capturing 90% of spectral density in the IMF with iterated sine in it. Spectral peaks from the iterated sine were removed from the data and linearly interpolated before calculating the signal-independent bandwidth. We found itEMD works to significantly increase the bandwidth of IMFs of interest, thereby capturing more waveform shape details (Figure S2). The effect is significant compared to existing sifting techniques across a wide range of noise conditions, but especially for the high-noise regime of σ>1. The bandwidth slightly increased with increasing frequency distortion, but only marginally. This may be because of the specific waveform used.

The drawback of higher bandwidth is capturing more noise together with signal. To evaluate this, we simulated 25s of zero-mean white noise with 10 cycles of 4Hz 8^th^ order iterated sine added to it. Noise was calculated as the standard deviation of 3s of signal away from the burst and signal was taken to be the mean instantaneous amplitude in the burst above the noise level. Signal-to-noise ratio (SNR, S/N) was computed as a summary metric against varying noise RMS. For high noise levels (σ>1.5), intermittent bursts in data were boosted less in SNR compared with ensemble and dyadic mask sifting due to higher noise coming from higher bandwidth. If non-sinusoidality is not of interest to the researcher and one is trying to identify a burst with very low SNR, other sifting techniques may be more suitable than itEMD. It is also pertinent to note that itEMD still has bandwidth limitations and cannot recover all of a harmonic which is at frequencies much higher to the original signal. For the iterated sine studied here, this cut-off was in the region where harmonic frequency is much above the base frequency, i.e. f_harmonic_>> 1.5 * f_base_. In general, this will depend on the amplitude ratios of the base and harmonic as well as their frequencies, but our results suggest itEMD treats signal separation similarly to original EMD with no noise (53).

**Figure S2:**
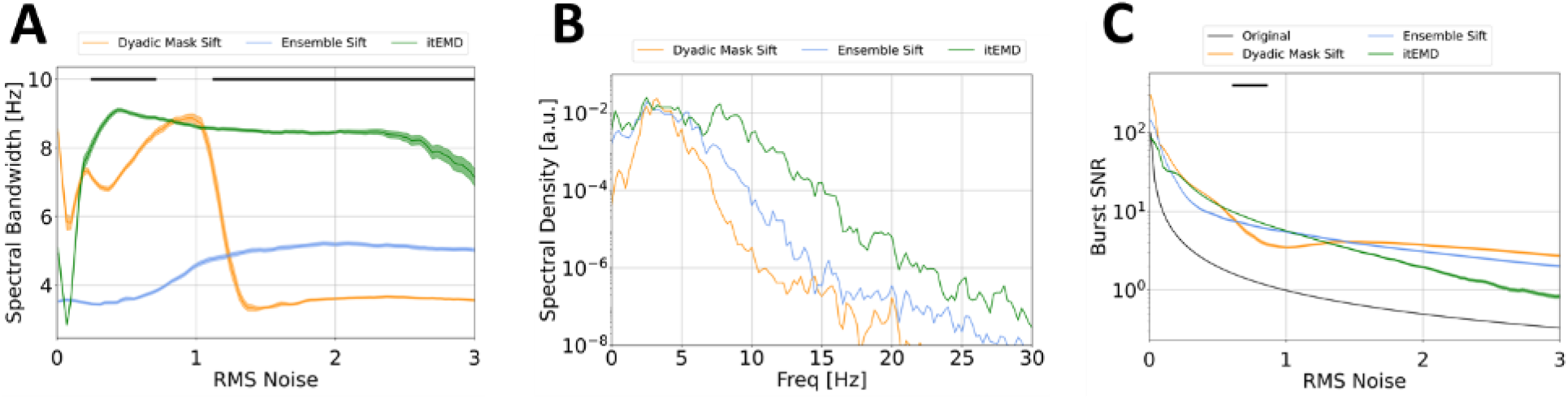
Bandwidth limitations of EMD. (A) Spectral bandwidth of signal IMF against white noise RMS. The novel itEMD is able to capture more waveform shape details due to significantly higher bandwidth. Black line indicates itEMD higher than the best existing method (P<0.01, Bonferroni-corrected). (B) Example signal IMF spectral density for RMS noise σ=1.5 (iterated sine spectral peaks removed for bandwidth calculation). (C) Signal-to-noise (SNR) improvement for an intermittent 4Hz burst in white noise of σ=1. The downside of higher bandwidth of itEMD is allowing more noise, which lowers SNR boost compared to existing techniques. All simulations were run with N=100 different noise realisations; mean ± standard error are shown.

### S3. Brown Noise Simulations

In the main text, we presented results for white noise simulations, as they are generalisable to signals with any frequency. Here we briefly present further results from a more neurophysiologically motivated set of simulations.

A variety of electrophysiological recordings across species and modalities has shown presence of fractal 1/f noise (also called aperiodic activity, scale-free activity, and pink / brown noise) (56–59). As such, we simulated ten seconds of brown noise (25s for sparsity simulations) with the colorednoise Python package (https://pypi.org/project/colorednoise/, based on Timmer and Koenig (1995) (60)). Data was sampled at 512Hz with low-frequency noise cut-off at 0.1Hz. To this 4Hz iterated sine signal was added, and quality of waveform shape reconstruction was evaluated with Pearson correlation to ground truth phase-aligned IF and the PMSI as in the main text. The same simulations exploring effect of noise, shape distortion, and sparsity were added (Figures S3). The results were very similar to those from white noise simulations. The novel itEMD outperformed existing sifting methods across a range of noise and shape distortion parameters, with the effect was most pronounced for highly noisy, highly non-sinusoidal signals. Iterated masking also significantly reduced mode mixing as measured by the PMSI (P<0.01, Bonferroni-corrected). Example sifting results can be seen in Figure S4. itEMD reconstructs the burst best of the three sifting methods compared and requires fewest IMFs to capture data completely due to higher IMF bandwidth, allowing for easier interpretation of modes and further analysis.

**Figure S3:**
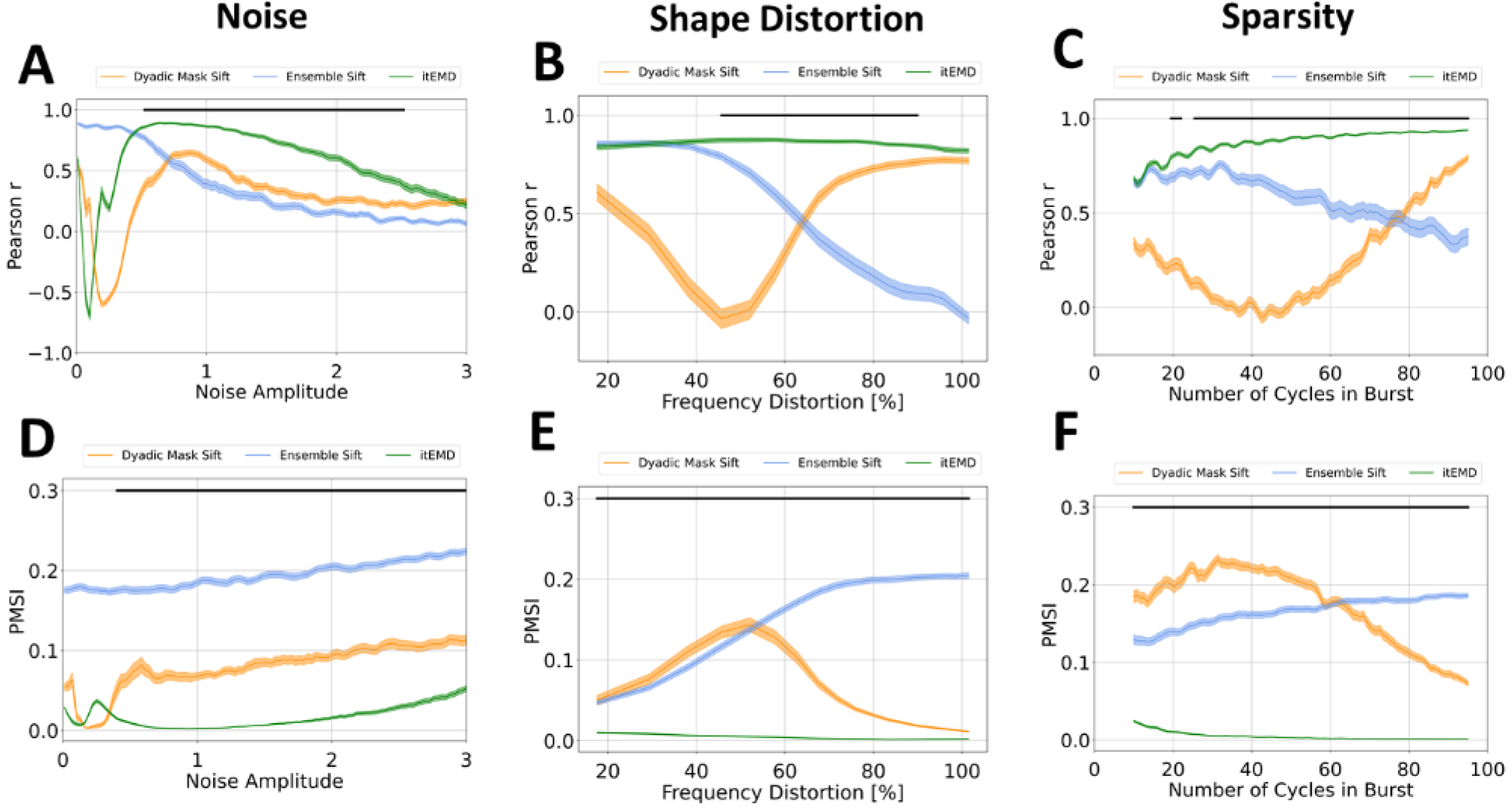
Brown noise simulation results. (A) Pearson correlation to ground truth waveform shape against brown noise amplitude (RMS). (B) Pearson r with ground truth against changing frequency distortion. (C) Pearson r against varying number of cycles in a sparse burst. (D) Pseudo-mode splitting index (PMSI) against brown noise amplitude. (E) PMSI against frequency distortion. (F) PMSI against number of cycles in a sparse burst. In all figures, black line indicates itEMD performing better (higher r, lower PMSI) than the best existing method (P<0.01, Bonferroni-corrected). itEMD can reconstruct waveform shape details for highly non-sinusoidal oscillations even in highly noisy sparse datasets.

**Figure S4:**
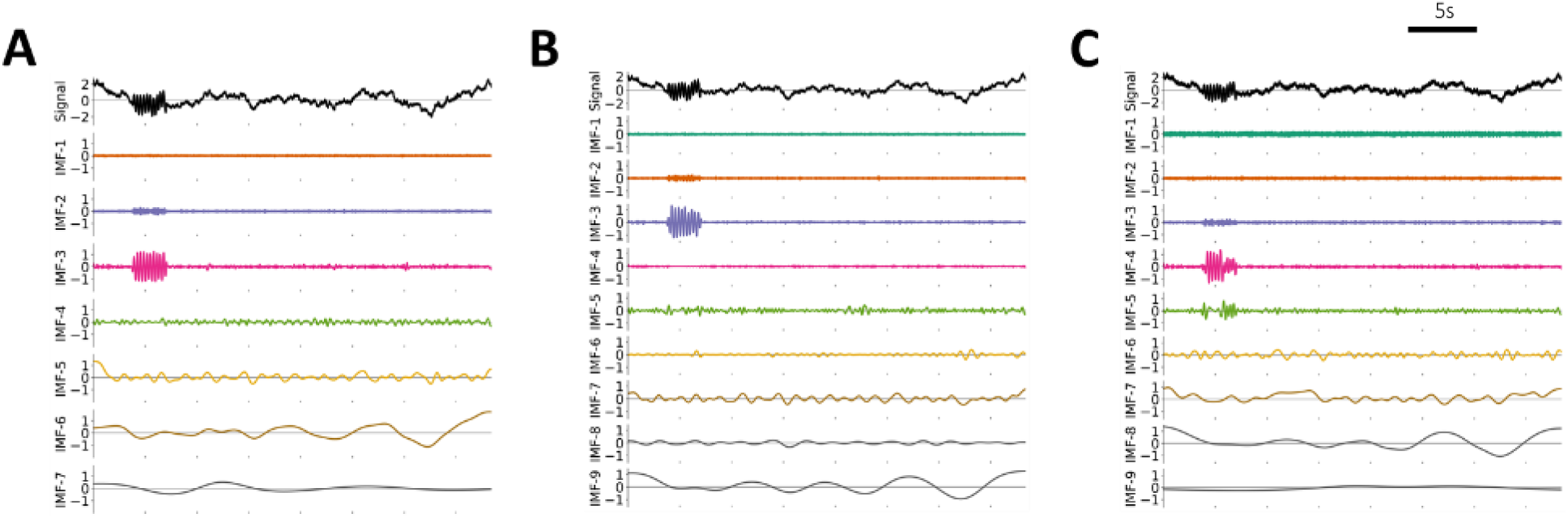
Example brown noise sifting results. IMFs are decompositions of 30 seconds of brown noise of amplitude 0.75 with a 10-cycle burst of 8^th^ order 4Hz iterated sine signal (top panels, “Signal”). (A) IMFs for the novel itEMD. Burst is well-reconstructed in IMF-3 and data is captured in 7 modes due to higher mode bandwidth. (B) IMFs for dyadic mask sifting. More mode mixing is present, details of shape are not resolved, and 9 IMFs are needed to capture data due to lower IMF bandwidth. (C) IMFs for ensemble EMD sifting. Severe mode mixing is present (burst split across IMF-4 and IMF-5) and shape details are not captured.

### S4. Mask frequency variability

As mentioned in the Results, IMFs known to contain signal had significantly lower mask frequency variability between iterations. We quantified this behaviour during the noise simulations described in the Main Text (Figure S5). At noise RMS σ=1.5, the relative difference between convergent mask frequencies of IMFs and those during the previous iteration were recorded. Based on the instantaneous frequency, signal was known to be in IMF-4. The variability of its mask frequency was compared to IMF-5, the below mode containing noise only. The mask frequency variability was significantly lower in the IMF known to have signal (P=1.01e-23). Noise IMF across N=100 noise realisations had a uniform range of mask variabilities under the iteration convergence threshold of 10%. Signal IMF variabilities were all under 3%, with most even lower. This shows itEMD robustly converges on signal when it is present. In the future, mask variability could thus be used to cross-validate the presence of true oscillations in an IMF.

**Figure S5:**
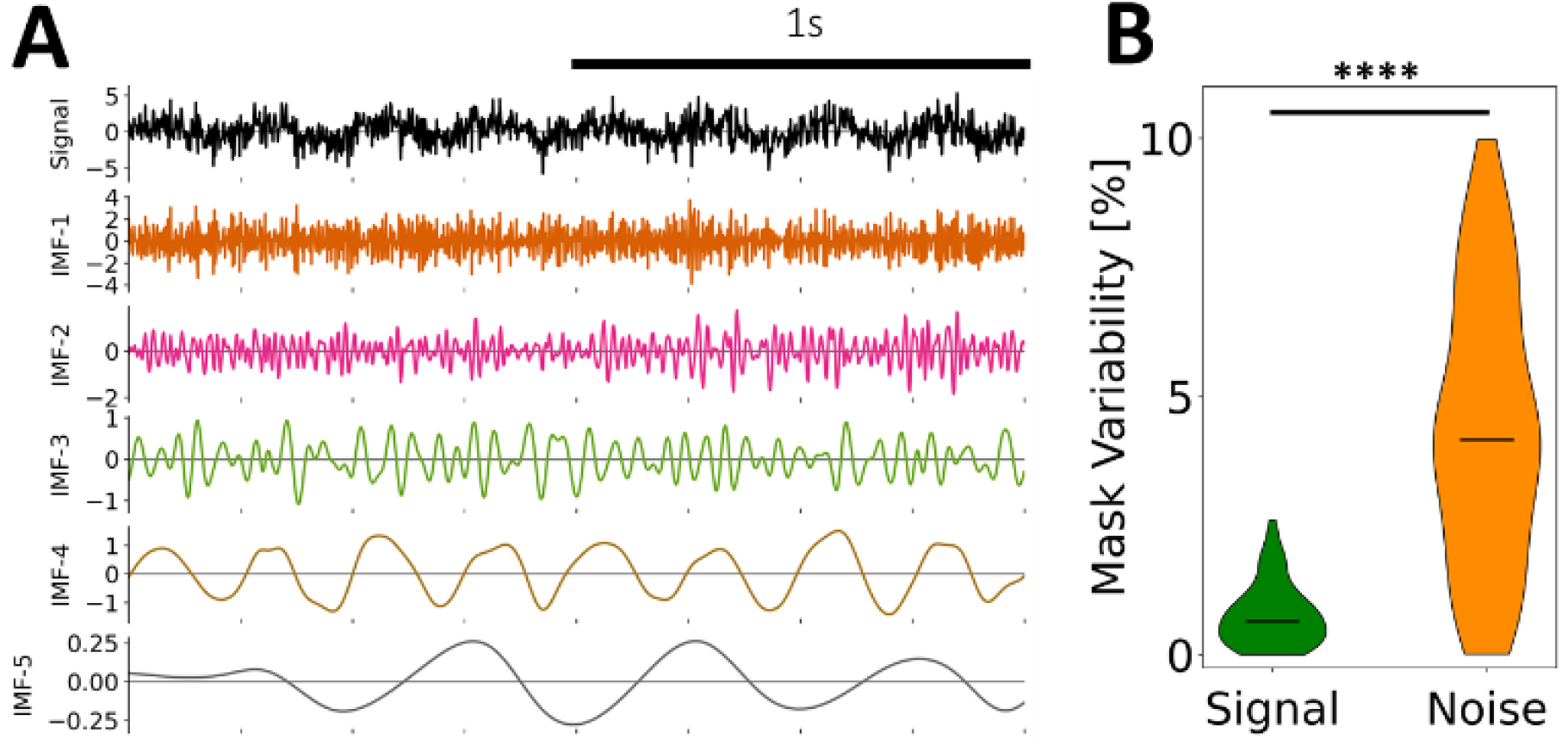
ItEMD has significantly less mask variability between iterations when a real signal is present. (A) Example simulated 2s of white noise and iterated sine signal. Signal is in IMF-3, noise in other IMFs. (B) Relative difference between convergent mask frequencies and those in the previous iteration. Mask is significantly less variable at convergence for IMF-4 containing signal compared to IMF-5 with noise only.

### S5. Single segment MEG results

As mentioned in the Discussion, the MEG analysis was re-run without segmentation of the data to verify alpha did not drift significantly throughout the recordings. The rest of the analysis of identical to that outlined in the Main Text. The single-segment results were very similar to results from segmenting recordings into 10 parts (Figure S5). Mode mixing measured by the PMSI was significantly reduced compared to existing sifting methods (P = 3.49e-5 vs dyadic mask, P = 0.0119 vs EEMD for lower PMSI, one-sided Bonferroni-corrected paired t-test), but was not significantly different to segmented results. Iterated masking also reproduced between-subject variance in mean occipital alpha frequency (P = 0.0113, F = 11.61, F-test against constant value). In summary, segmenting recordings may improve performance, but yields very similar results when shape is not changing significantly.

**Figure S6:**
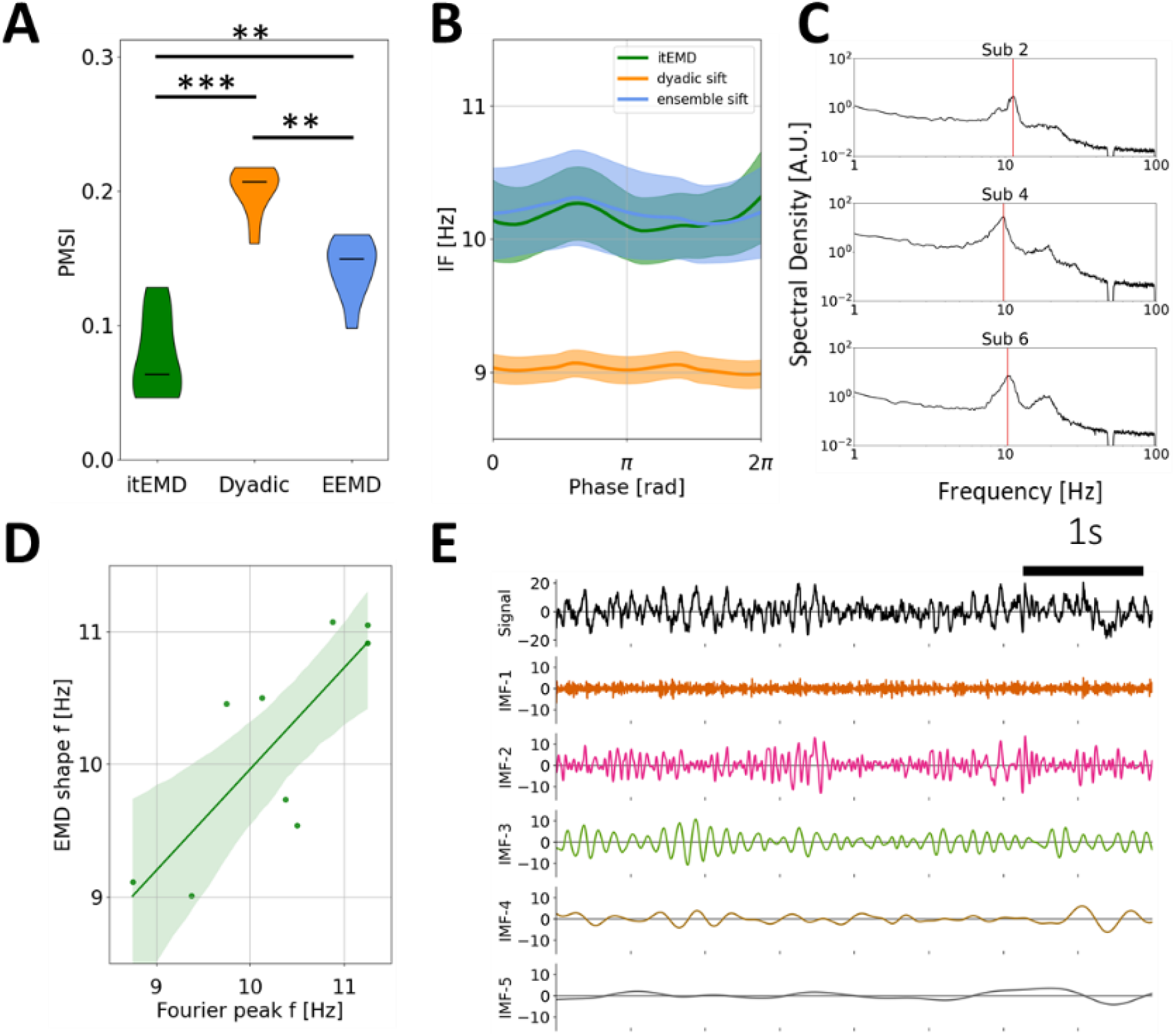
Human MEG occipital alpha results without data segmentation is similar to segmented results in Main Text. (A) Group-level violin plots of the pseudo-mode splitting index (PMSI, a measure of mode mixing). Iterated masking had significantly lower PMSI than both dyadic mask sift (P = 3.49e-5) and ensemble sift (P = 0.0119, both Bonferroni-corrected). (B) Group-level phase-aligned instantaneous frequency (mean± standard error shaded). Both itEMD and EEMD detect the 10Hz occipital alpha oscillation with no significant non-sinusoidality. Masked sift fails to capture the oscillation well due to mode mixing. (C) Example Fourier power spectra for individual subjects. Alpha peak is denoted with the red line. Note the presence of a second beta-band peak. (D) Mean subject phase-aligned instantaneous frequency against peak alpha frequency from the Fourier power spectral density. Iterated masking linearly reproduces between-subject variability in alpha frequency (P=0.0113, F-test against constant null hypothesis). (E) Example 5s of raw itEMD sift results. Alpha oscillations are in IMF-3 with sharp features in IMF-2 and minimal mode mixing.

## REFERENCES

1. Cole SR, Voytek B. Brain Oscillations and the Importance of Waveform Shape. Trends Cogn Sci 21: 137–149, 2017. doi: 10.1016/j.tics.2016.12.008.

2. Hari R, Puce A. MEG-EEG primer. New York, NY: Oxford University Press, 2017.

3. Lo M-T, Novak V, Peng C-K, Liu Y, Hu K. Nonlinear phase interaction between nonstationary signals: A comparison study of methods based on Hilbert-Huang and Fourier transforms. Phys Rev E Stat Nonlin Soft Matter Phys 79: 061924, 2009. doi: 10.1103/PhysRevE.79.061924.

4. Lozano-Soldevilla D, ter Huurne N, Oostenveld R. Neuronal Oscillations with Non-sinusoidal Morphology Produce Spurious Phase-to-Amplitude Coupling and Directionality. Front Comput Neurosci 10, 2016. doi: 10.3389/fncom.2016.00087.

5. Vidaurre D, Quinn AJ, Baker AP, Dupret D, Tejero-Cantero A, Woolrich MW. Spectrally resolved fast transient brain states in electrophysiological data. NeuroImage 126: 81–95, 2016. doi: 10.1016/j.neuroimage.2015.11.047.

6. Bartz S, Avarvand FS, Leicht G, Nolte G. Analyzing the waveshape of brain oscillations with bicoherence. NeuroImage 188: 145–160, 2019. doi: 10.1016/j.neuroimage.2018.11.045.

7. Amzica F, Steriade M. Electrophysiological correlates of sleep delta waves. Electroencephalogr Clin Neurophysiol 107: 69–83, 1998. doi: 10.1016/S0013-4694(98)00051-0.

8. Belluscio MA, Mizuseki K, Schmidt R, Kempter R, Buzsáki G. Cross-Frequency Phase–Phase Coupling between Theta and Gamma Oscillations in the Hippocampus. J Neurosci 32: 423–435, 2012. doi: 10.1523/JNEUROSCI.4122-11.2012.

9. Cole SR, Meij R van der, Peterson EJ, Hemptinne C de, Starr PA, Voytek B. Nonsinusoidal Beta Oscillations Reflect Cortical Pathophysiology in Parkinson’s Disease. J Neurosci 37: 4830–4840, 2017. doi: 10.1523/JNEUROSCI.2208-16.2017.

10. Huang NE, Shen Z, Long SR, Wu MC, Shih HH, Zheng Q, Yen N-C, Tung CC, Liu HH. The empirical mode decomposition and the Hilbert spectrum for nonlinear and non-stationary time series analysis. Proc R Soc Lond Ser Math Phys Eng Sci 454: 903–995, 1998. doi: 10.1098/rspa.1998.0193.

11. Quinn AJ, Lopes-dos-Santos V, Huang N, Liang W-K, Juan C-H, Yeh J-R, Nobre AC, Dupret D, Woolrich MW. Within-cycle instantaneous frequency profiles report oscillatory waveform dynamics. bioRxiv, 2021. doi: 10.1101/2021.04.12.439547v1

12. Giehl J, Noury N, Siegel M. Dissociating harmonic and non-harmonic phase-amplitude coupling in the human brain. NeuroImage 227: 117648, 2021. doi: 10.1016/j.neuroimage.2020.117648.

13. Yang Y, Deng J, Wu C. Analysis of Mode Mixing Phenomenon in the Empirical Mode Decomposition Method. In: 2009 Second International Symposium on Information Science and Engineering. 2009 International Symposium on Information Science and Engineering (ISISE). IEEE, p. 553–556.

14. Bueno-López M, Giraldo E, Molinas M, Fosso OB. The Mode Mixing Problem and its Influence in the Neural Activity Reconstruction..

15. Lundqvist M, Rose J, Herman P, Brincat SL, Buschman TJ, Miller EK. Gamma and Beta Bursts Underlie Working Memory. Neuron 90: 152–164, 2016. doi: 10.1016/j.neuron.2016.02.028.

16. Feingold J, Gibson DJ, DePasquale B, Graybiel AM. Bursts of beta oscillation differentiate postperformance activity in the striatum and motor cortex of monkeys performing movement tasks. Proc Natl Acad Sci 112: 13687–13692, 2015. doi: 10.1073/pnas.1517629112.

17. van Ede F, Quinn AJ, Woolrich MW, Nobre AC. Neural Oscillations: Sustained Rhythms or Transient Burst-Events? Trends Neurosci 41: 415–417, 2018. doi: 10.1016/j.tins.2018.04.004.

18. Wu Z, Huang NE. A study of the characteristics of white noise using the empirical mode decomposition method. Proc R Soc Lond Ser Math Phys Eng Sci 460: 1597–1611, 2004. doi: 10.1098/rspa.2003.1221.

19. Flandrin P, Rilling G, Goncalves P. Empirical mode decomposition as a filter bank. IEEE Signal Process Lett 11: 112–114, 2004. doi: 10.1109/LSP.2003.821662.

20. Wu Z, Huang NE. Ensemble empirical mode decomposition: a noise-assisted data analysis method. Adv Adapt Data Anal 01: 1–41, 2009. doi: 10.1142/S1793536909000047.

21. Deering R, Kaiser JF. The use of a masking signal to improve empirical mode decomposition. In: Proceedings. (ICASSP ’05). IEEE International Conference on Acoustics, Speech, and Signal Processing, 2005. Proceedings. (ICASSP ’05). IEEE International Conference on Acoustics, Speech, and Signal Processing, 2005. 2005, p. iv/485–iv/488 Vol. 4.

22. Wang Y-H, Hu K, Lo M-T. Uniform Phase Empirical Mode Decomposition: An Optimal Hybridization of Masking Signal and Ensemble Approaches. IEEE Access 6: 34819–34833, 2018. doi: 10.1109/ACCESS.2018.2847634.

23. Wang H, Ji Y. A Revised Hilbert–Huang Transform and Its Application to Fault Diagnosis in a Rotor System. Sensors 18: 4329, 2018. doi: 10.3390/s18124329.

24. Yang Y, Deng J, Kang D. An improved empirical mode decomposition by using dyadic masking signals. Signal Image Video Process 9: 1259–1263, 2015. doi: 10.1007/s11760-013-0566-7.

25. Fosso OB, Molinas M. Method for Mode Mixing Separation in Empirical Mode Decomposition [Online]. http://arxiv.org/abs/1709.05547 [20 Jan. 2021].

26. Torres ME, Colominas MA, Schlotthauer G, Flandrin P. A complete ensemble empirical mode decomposition with adaptive noise. In: 2011 IEEE International Conference on Acoustics, Speech and Signal Processing (ICASSP). ICASSP 2011 - 2011 IEEE International Conference on Acoustics, Speech and Signal Processing (ICASSP). IEEE, p. 4144–4147.

27. Yeh J-R, Shieh J-S, Huang NE. Complementary ensemble empirical mode decomposition: a novel noise enhanced data analysis method. Adv Adapt Data Anal 02: 135–156, 2010. doi: 10.1142/S1793536910000422.

28. Shafto MA, Tyler LK, Dixon M, Taylor JR, Rowe JB, Cusack R, Calder AJ, Marslen-Wilson WD, Duncan J, Dalgleish T, Henson RN, Brayne C, Matthews FE, Cam-CAN. The Cambridge Centre for Ageing and Neuroscience (Cam-CAN) study protocol: a cross-sectional, lifespan, multidisciplinary examination of healthy cognitive ageing. BMC Neurol 14: 204, 2014. doi: 10.1186/s12883-014-0204-1.

29. Taylor JR, Williams N, Cusack R, Auer T, Shafto MA, Dixon M, Tyler LK, Cam-CAN, Henson RN. The Cambridge Centre for Ageing and Neuroscience (Cam-CAN) data repository: Structural and functional MRI, MEG, and cognitive data from a cross-sectional adult lifespan sample. NeuroImage 144: 262–269, 2017. doi: 10.1016/j.neuroimage.2015.09.018.

30. Quinn AJ, Lopes-dos-Santos V, Dupret D, Nobre AC, Woolrich MW. EMD: Empirical Mode Decomposition and Hilbert-Huang Spectral Analyses in Python. J Open Source Softw 6: 2977, 2021. doi: 10.21105/joss.02977.

31. Harris CR, Millman KJ, van der Walt SJ, Gommers R, Virtanen P, Cournapeau D, Wieser E, Taylor J, Berg S, Smith NJ, Kern R, Picus M, Hoyer S, van Kerkwijk MH, Brett M, Haldane A, del Río JF, Wiebe M, Peterson P, Gérard-Marchant P, Sheppard K, Reddy T, Weckesser W, Abbasi H, Gohlke C, Oliphant TE. Array programming with NumPy. Nature 585: 357–362, 2020. doi: 10.1038/s41586-020-2649-2.

32. Virtanen P, Gommers R, Oliphant TE, Haberland M, Reddy T, Cournapeau D, Burovski E, Peterson P, Weckesser W, Bright J, van der Walt SJ, Brett M, Wilson J, Millman KJ, Mayorov N, Nelson ARJ, Jones E, Kern R, Larson E, Carey CJ, Polat İ, Feng Y, Moore EW, VanderPlas J, Laxalde D, Perktold J, Cimrman R, Henriksen I, Quintero EA, Harris CR, Archibald AM, Ribeiro AH, Pedregosa F, van Mulbregt P. SciPy 1.0: fundamental algorithms for scientific computing in Python. Nat Methods 17: 261–272, 2020. doi: 10.1038/s41592-019-0686-2.

33. Seabold S, Perktold J. Statsmodels: Econometric and Statistical Modeling with Python. Python in Science Conference, p. 92–96.

34. Hunter JD. Matplotlib: A 2D Graphics Environment. Comput Sci Eng 9: 90–95, 2007. doi: 10.1109/MCSE.2007.55.

35. Cole S, Voytek B. Cycle-by-cycle analysis of neural oscillations. J Neurophysiol 122: 849–861, 2019. doi: 10.1152/jn.00273.2019.

36. Sherin B. Common sense clarified: The role of intuitive knowledge in physics problem solving. J Res Sci Teach 43: 535–555, 2006. doi: 10.1002/tea.20136.

37. Boashash B. Estimating and interpreting the instantaneous frequency of a signal. I. Fundamentals. Proc IEEE 80: 520–538, 1992. doi: 10.1109/5.135376.

38. Huang NE, Wu Z, Long SR, Arnold KC, Chen X, Blank K. On instantaneous frequency. Adv Adapt Data Anal 01: 177–229, 2009. doi: 10.1142/S1793536909000096.

39. Alabdulmohsin IM. Theorems and Methods on Partial Functional Iteration. Proceedings of the World Congress on Engineering, 2009.

40. Cranstoun SD, Ombao HC, von Sachs R, Wensheng Guo, Litt B. Time-frequency spectral estimation of multichannel EEG using the Auto-SLEX method. IEEE Trans Biomed Eng 49: 988–996, 2002. doi: 10.1109/TBME.2002.802015.

41. Salmelin R, Hari R. Spatiotemporal characteristics of sensorimotor neuromagnetic rhythms related to thumb movement. Neuroscience 60: 537–550, 1994. doi: 10.1016/0306-4522(94)90263-1.

42. Sedgwick P. Multiple significance tests: the Bonferroni correction. BMJ 344: e509, 2012. doi: 10.1136/bmj.e509.

43. Mizuseki K, Diba K, Pastalkova E, Teeters J, Sirota A, Buzsáki G. Neurosharing: large-scale data sets (spike, LFP) recorded from the hippocampal-entorhinal system in behaving rats. F1000Research 3, 2014. doi: 10.12688/f1000research.3895.1.

44. Mizuseki K, Sirota A, Pastalkova E, Diba K, Buzsáki G. Multiple single unit recordings from different rat hippocampal and entorhinal regions while the animals were performing multiple behavioral tasks. CRCNS.org [Online]. 2013. http://dx.doi.org/10.6080/K09G5JRZ [29 Mar. 2021].

45. Donoghue T, Haller M, Peterson EJ, Varma P, Sebastian P, Gao R, Noto T, Lara AH, Wallis JD, Knight RT, Shestyuk A, Voytek B. Parameterizing neural power spectra into periodic and aperiodic components. Nat Neurosci 23: 1655–1665, 2020. doi: 10.1038/s41593-020-00744-x.

46. Buzsáki G, Czopf J, Kondákor I, Kellényi L. Laminar distribution of hippocampal rhythmic slow activity (RSA) in the behaving rat: Current-source density analysis, effects of urethane and atropine. Brain Res 365: 125–137, 1986. doi: 10.1016/0006-8993(86)90729-8.

47. Buzsáki G, Rappelsberger P, Kellényi L. Depth profiles of hippocampal rhythmic slow activity (’theta rhythm’) depend on behaviour. Electroencephalogr Clin Neurophysiol 61: 77–88, 1985. doi: 10.1016/0013-4694(85)91075-2.

48. Haegens S, Cousijn H, Wallis G, Harrison PJ, Nobre AC. Inter- and intra-individual variability in alpha peak frequency. NeuroImage 92: 46–55, 2014. doi: 10.1016/j.neuroimage.2014.01.049.

49. Klimesch W. EEG-alpha rhythms and memory processes. Int J Psychophysiol 26: 319–340, 1997. doi: 10.1016/S0167-8760(97)00773-3.

50. Siapas AG, Lubenov EV, Wilson MA. Prefrontal Phase Locking to Hippocampal Theta Oscillations. Neuron 46: 141–151, 2005. doi: 10.1016/j.neuron.2005.02.028.

51. Gips B, Bahramisharif A, Lowet E, Roberts MJ, de Weerd P, Jensen O, van der Eerden J. Discovering recurring patterns in electrophysiological recordings. J Neurosci Methods 275: 66–79, 2017. doi: 10.1016/j.jneumeth.2016.11.001.

52. Jas M, La Tour TD, Şimşekli U, Gramfort A. Learning the Morphology of Brain Signals Using Alpha-Stable Convolutional Sparse Coding [Online]. http://arxiv.org/abs/1705.08006 [29 Mar. 2021].

53. Rilling G, Flandrin P. One or Two Frequencies? The Empirical Mode Decomposition Answers. IEEE Trans Signal Process 56: 85–95, 2008. doi: 10.1109/TSP.2007.906771.

54. Aru J, Aru J, Priesemann V, Wibral M, Lana L, Pipa G, Singer W, Vicente R. Untangling cross-frequency coupling in neuroscience. Curr Opin Neurobiol 31: 51–61, 2015. doi: 10.1016/j.conb.2014.08.002.

55. Gerber EM, Sadeh B, Ward A, Knight RT, Deouell LY. Non-Sinusoidal Activity Can Produce Cross-Frequency Coupling in Cortical Signals in the Absence of Functional Interaction between Neural Sources. PLOS ONE 11: e0167351, 2016. doi: 10.1371/journal.pone.0167351.

56. He BJ, Zempel JM, Snyder AZ, Raichle ME. The Temporal Structures and Functional Significance of Scale-free Brain Activity. Neuron 66: 353–369, 2010. doi: 10.1016/j.neuron.2010.04.020.

57. Voytek B, Knight RT. Dynamic Network Communication as a Unifying Neural Basis for Cognition, Development, Aging, and Disease. Biol Psychiatry 77: 1089–1097, 2015. doi: 10.1016/j.biopsych.2015.04.016.

58. Miller KJ, Sorensen LB, Ojemann JG, Nijs M den. Power-Law Scaling in the Brain Surface Electric Potential. PLOS Comput Biol 5: e1000609, 2009. doi: 10.1371/journal.pcbi.1000609.

59. Colombo MA, Napolitani M, Boly M, Gosseries O, Casarotto S, Rosanova M, Brichant J-F, Boveroux P, Rex S, Laureys S, Massimini M, Chieregato A, Sarasso S. The spectral exponent of the resting EEG indexes the presence of consciousness during unresponsiveness induced by propofol, xenon, and ketamine. NeuroImage 189: 631–644, 2019. doi: 10.1016/j.neuroimage.2019.01.024.

60. Timmer J, König M. On Generating Power Law Noise. A&a 300: 707–710, 1995.

